# Lossless Single-Molecule Counting To Absolute Quantify Proteoforms

**DOI:** 10.1101/2024.03.19.585761

**Authors:** Tobias Gross, Tobias Hundertmark, Villő Csiszár, András Attila Sulyok, Nina Gross, Maike Breiden, Niklas Kitschen, Uritza von Groll, Christoph Niemöller, Pablo Sánchez-Martín, Anne Hein, Jens Göpffert, Tamás Szórádi, Philipp Lübbert, Peter Koltay, Peter Porschewski, Felix von Stetten, Roland Zengerle, Csaba Jeney

**Affiliations:** Laboratory for MEMS Applications, Department of Microsystems Engineering, University of Freiburg, Georges-Köhler-Allee 103, Freiburg, 79110, Germany; Actome GmbH, Georges-Köhler-Allee 103, Freiburg, 79110, Germany; Faculty of Information Technology and Bionics, Pázmány Péter Catholic University, Práter Street 50/A, Budapest, Hungary; Department of Probability Theory and Statistics, Institute of Mathematics, Eötvös Loránd University, Pázmany Péter sétány 1/C, Budapest, 1117, Hungary; Hahn-Schickard, Georges-Köhler-Allee 302, Freiburg, 79110, Germany; QIAGEN, Qiagen Strasse 1, Hilden, 40724, Germany; NMI, Markwiesenstraße 55, Reutlingen, Germany

## Abstract

A novel immunoassay, termed Protein Interaction Coupling (PICO), is introduced to deliver unequivocal, reference-free quantification of proteoforms - absolute quantification. PICO employs a compartmentalized, homogeneous single-molecule assay with a lossless and highly sensitive signal generation, capable of detecting down to a few molecules per reaction. Additionally, it utilises a background-free, digital enumeration principle, known as the decouplexing. PICO is presented as exact mathematical theories, providing a theoretical comprehension of its chemistry. Consequently, PICO demonstrates absolute quantification, as exemplified with recombinant and non-recombinant ErbB2 and multi-tagged peptide rTRX targets, validating absolute quantification against internal and external references in both analytical and cellular matrices. Furthermore, PICO enables combinatorial multiplexing (cplex), a readout between any two antibodies, demonstrated by an 8-plex antibody, 12-cplex PICO, measuring functional changes of ErbB pathway upon mock and dactolisib treatment delivering absolute quantitative cellular stoichiometry. PICO possesses immense potential for versatile, standardized, and accurate protein measurements, offering insights into physiological and perturbed cellular processes.

---

The labyrinthine complexity of the human genome is a marvel to behold. The expressed proteome is a realm of staggering complexity, boasting an estimated one million proteoforms [1]. A particularly salient chemical attribute is the dynamic concentration of proteoforms including proteins, PPIs, PTMs and overall material (and information) flows through the networks, as these serve as the primary regulated chemical actors related to the physiology of living cell. Monitoring the proteoforms has been a major aim of protein analytical methods [2], however, a significant challenge that persists is the accurate determination of absolute protein concentrations with universal comparability between measurements especially between different proteoforms.

Generally, absolute quantitative protein assays require a readout method ensuring there is no off-setting background signal, i.e. having zero background, and that the generated quantitative signal is robust, capable of providing lossless molecular counts in a wide and linear dynamic range, i.e. absolute signal. These are seemingly conflicting requirements due to the required extreme signal-to-noise performance. Due to these hurdles, absolute quantificative analytical methods are only partially established in the study of biological systems, and mainly based on mass spectrometry techniques [3]. Methods without using external quantitative standards, avoiding unreliabilities sourced form the vastly different qualities of quantitative protein standards, are especially scarce. Specifically for immunoassays a reference-free absolute quantification is not yet established despite the significant advance in digital immunoassays, exemplifying the most advanced technologies including SIMOA and others [4, 5].

## Results

### Lossless, zero background detection principle - partitioning of single molecules, analytical limit of detection

We have pioneered an approach, termed *protein interaction coupling* (PICO [6, 7]), explicitly architected to transcend existing methods (for workflow Fig 1.a). PICO achieves absolute, lossless quantification of tar-get proteoforms through (1) the single molecule sensitivity of the applied compartmentalized PCR detection and the physically compartmentalized signal generation (dPCR). Additionally (2) the absence of background-contributing side reactions guarantees (3) zero background signal borrowed from the absolute, digital counting of dPCR. Moreover, it is (4) independent of antibody dissociation constants (*K*_*d*_) by choosing carefully orchestrated chemical conditions, and importantly, PICO is (5) homogeneous ensuring a robust, lossless process, without washing steps. The (6) high specificity of PICO is defined by two antibodies conjugated to amplifiable DNA labels, strictly binding of both asserts the signal. Thereby drawing conceptual parallels to other bi-component assays that are predominantly enzyme-label reliant, however, in the case of PICO, (7) under stringent, aspecific binding suppressive conditions. The PICO readout are not being restricted to given, cognate antibody pair, as signals between any antibody pairs (ABP) can be read out. This is termed (8) combinatorial multiplexing (cplex), exemplified here as triangular, quadratic (three and six ABPs). The number of ABPs that can be analyzed has recently reached to 12, with the potential to reach at least a hundred in the foreseeable future. For the sake of simplicity the term *couplex* is defined as the ternary complex constituted by the two antibodies and the target proteoform. The couplexes are quantified (see Fig 1.a for general dPCR concept see [8]) by a novel statistical model tailored to analyze data based on Poisson-distribution [9] called *compartment decouplexing*, or DX, (for the model see Appendix A). Utilizing this model, three pivotal parameters (concentration of couplexes, and of the two antibodies) are extracted that comprehensively characterize the individual reactions to derive the absolute couplex signal. Consequently, PICO readout manifests an outstanding analytical performance, as corroborated by the exceedingly low limit of absolute detection of DNA-based artificial couplexes (LOD=3 molecules per reaction) illustrated in Fig 1.b. In addition to points (2) and (3), any antibody co-localizations, which are not a couplexes, are efficiently filtered out by the decouplexing model, which decisively contributes to the digital enumeration of the signal. This clearly illustrated in Fig 1.b displaying a unit of slope and a Y-intercept at zero. However, it should be noted that the derived couplexes are probabilistic variables with inherent calculable variance, as detailed in Appendix A and attested by simulation data (Fig A1), which in turn influences the precision of the assay. Also, couplex values are normally distributed (Appendix A).

**Figure 1:**
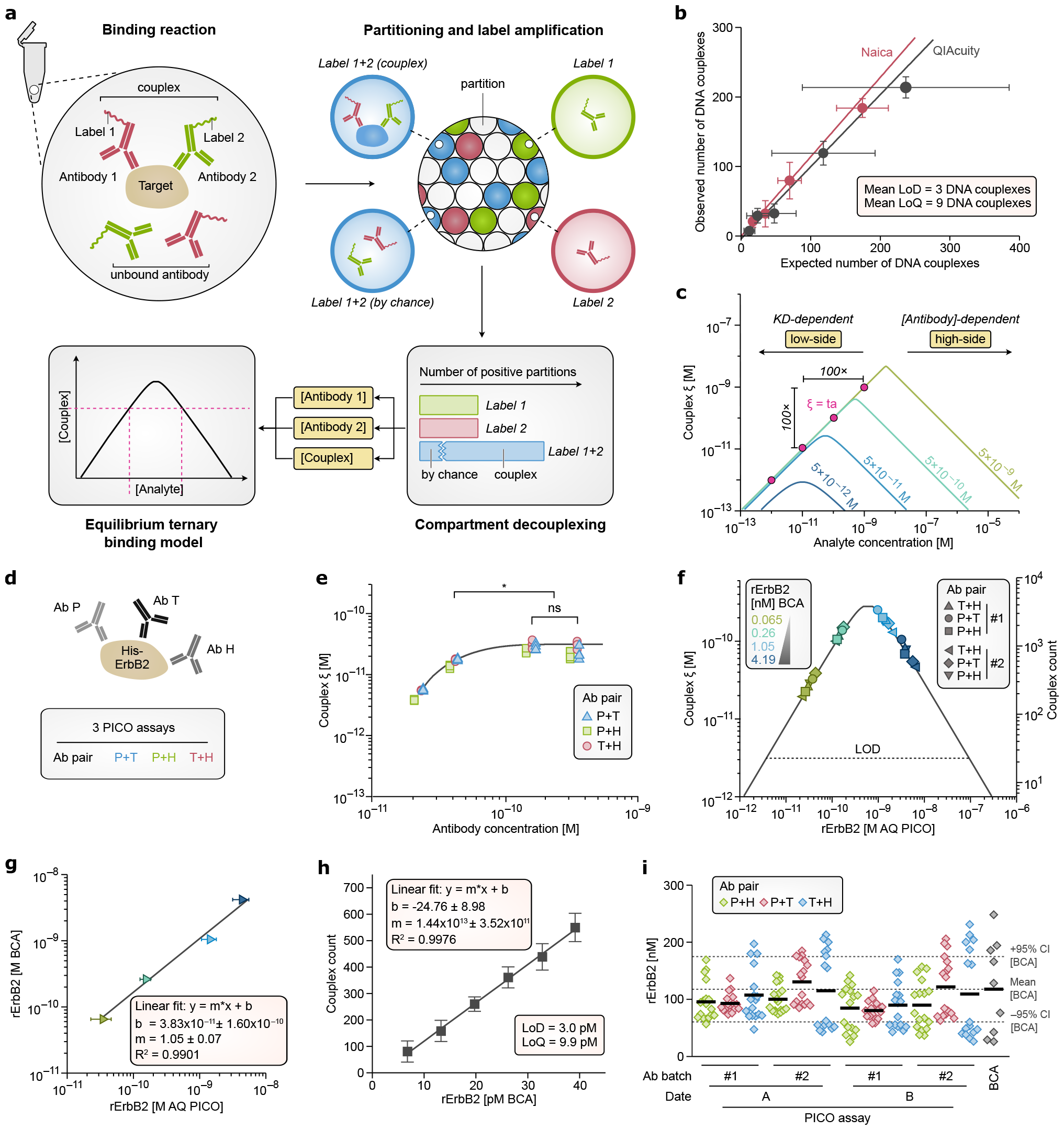
**a**. The schematic workflow for absolute quantification. The PICO binding reaction described by the AQ model (Appendix B), and consists of the analyte and two amplifiable DNA label-conjugated antibodies under appropriate binding concentrations and conditions (D.5). After incubation the reaction is subsequently diluted according to the conditions required for a single-molecule digital PCR (dPCR)D.4. To provide inputs for the AQ model, the concentrations of couplexes and antibodies required are determined using the decouplexing model (Section A), ultimately yielding the absolute concentration of analyte. Additionally, the labeling efficiency is considered for antibody concentration correction (Section D.3). **b**. Analytical PICO. Spiked-in versus observed number of artificial DNA couplexes applying the decouplexing model. The experimental details are in Method and Reagents D.6. The experiments were repeated on different devices as indicated. *R*^2^ is larger than 0.99 in all cases, the slopes are 1.04 ± 0.39 for Qiacuity and 1.14 ± 0.15 for Stilla, respectively, indicating 1-to-1 correspondence between spiked-in and measured amounts, showing lossless single-molecule counting. The average Y-intercept is −2.28 ± 10.21 couplex, indicating zero background. The mean analytical LOD of analytical PICO is 3 couplexes and the LOQ is 9 couplexes, based on the devised calculations of Wenzl et al. [28]. **c**. The non-bijective calibration curve of the AQ model relates absolute analyte concentration (x-axis) to measured couplex molar concentration (*ξ*, y-axis) in the binding reaction. It demonstrates the influence of decreasing antibody concentrations. Under saturated conditions, the three highest antibody concentrations, the *K*_*d*_-dependence of *ξ* is eliminated, as the *ξ* matches the target analyte concentration on the low side, while on the high side, the analyte concentration is calculated from the antibody concentrations using the AQ model. See AQ model in Section B for more details. **d**. Proof of absolute quantification. BCA verified amount of recombinant ErbB2, as quantitative external reference, was assayed in a triangular PICO setup using trastuzumab (T), pertuzumab (P) and anti-HIS (H) antibodies (see Materials and Methods D.1 and D.5). In the triangular PICO concept, the three antibodies forming three ABPs, which quantitative, pair-wise PICO AQ results must equal (internal verification) if absolute quantitativity holds. **e**. An isomolar (antigen) titration (D.5) was conducted at various concentrations of antibodies (13.3 pM, 40.0 pM, 120 pM, and 360 pM) with a fixed rErbB2 concentration of 40 pM. At the higher antibody concentrations, the generated number of couplexes did not significantly differ (Kruskal-Wallis ANOVA with Tukey’s test, *p <* 0.05), indicating that a saturation concentration of the standard ABX = 500 pM suffices for all antibodies. **f**. AQ (calibration) curve of recombinant ErbB2 (BCA verified). Symbols indicate different ABPs. The verified mean concentration of (all three) antibodies from the first batch is 5.29 × 10^−10^ ± 9.16 × 10^−11^ M (n=138) and for the second batch 5.66 × 10^−10^ ± 6.04 × 10^−11^ M (n=138), saturating concentration. The assay carried out using two different batches of antibodies (1 and 2), each marker representing the mean of 6 replicates and with standard deviations are shown (mainly imperceptible), n=216. The dotted line denotes the LOD of measurement (20 couplexes, LOD 3.00 pM). **g**. BCA reference vs. AQ PICO (data from Fig 1.c see also c.). A regression line has a slope of 1.05 ± 0.07, intercept 38.3 ± 160 pM and *R*^2^ = 0.99 with *n* = 216, confirms the linearity of the AQ model involving both sides of the curve. **h**. The relationship between the measured number of couplexes and the reference (BCA) values for the rErbB2 analyte, within close vicinity of the limit of detection (LOD), at 6.75 pM, 13.3 pM, 19.7 pM, 26.2 pM, 32.8 pM and 39.1 pM of rErbB2, respectively. LOD was 3.00 pM and LOQ 9.90 pM. Slope m = 14.4 ± 0.35, y-axis intersection *b* = −24.76 ± 8.98, *R*^2^ = 0.99, *n* = 36. **i**. Distribution of the measured absolute amounts of the rErbB2 quantitative reference normalised for dilution (see also c.). Two different time point, two different batches of antibodies, three ABPs and four dilution of of the analyte. The BCA quantified reference indicated (118 ± 87.7 nM) with 95 confidence intervals. Values from left to right are 95.44 ± 33.70 nM, 92.87 ± 16.60 nM, 107.28 ± 45.59 nM, 99.99 ± 23.62 nM, 130.52 ± 38.18 nM, 115.09 ± 69.39 nM, 84.65 ± 39.45 nM, 80.41 ± 15.41 nM, 89.77 ± 40.57 nM, 89.77 ± 40.57 nM, 121.64 ± 52.19 nM and 109.23 ± 79.41 nM stock concentration, respectively. ANOVA shows no significant differences observed in terms of ABPs, assay dates, and PICO compared to BCA (F(12,202) = 1.77, p-value = 0.0553, *n* = 211). These findings substantiate the absolute quantitative measurement of recombinant ErbB2, demonstrating lossless molecular counting, negligible background, and pronounced linearity over 3.5 log dynamic range.

### Theory of affinity of antibody independent absolute quantification

To derive the antibody *K*_*d*_-independent absolute quantification of proteoform targets, we have formulated and analyzed an *ternary-equilibrium-based absolute quantification model* (i.e. AQ model) (Appendix B, see for numerical analysis B2). We derived motivation from a range of models that have been utilized to study ternary binding in previous studies [10] (see other references in Appendix B), however our analysis is specific for solution phase ternary immunoassays and reveals implications for pioneering absolute quantitative measurements. According to analyses of AQ model in Appendix B, also see Fig 1.c, the model characterized as a non-bijective curve predicting two antigen concentrations for each measured couplex value. It is noteworthy that the dual readout effectively doubles the dynamic range achieving a dynamic range of five magnitudes routinely (Fig 1.f-g). At saturating concentrations of antibodies (Fig 1.e), according to the analysis the low side of the curve is negligibly influenced by the dissociation constant of antibodies (*K*_*d*_), whereas the high side is exclusively dependent on antibody concentration, devoid of *K*_*d*_ influence (Fig B4.a-b). The model further exhibits a slope of unity (and minus unity) as a dual linear behavior in log-log representation, thereby enabling, additionally, facile ratio calculations for relative couplex quantification (Fig 1.c). Consequently, *K*_*d*_-independent behavior in PICO assays is attainable at saturating antibody concentrations (Fig B5). Under these conditions, the concentration of couplexes equals with the concentration of the antigen on the low side, while the antigen concentration on the high-side can be directly inferred from the concentrations of antibodies and couplexes. In practical terms, the antigen concentrations of the model for both the low and high sides can be computed using an appropriate surrogate *K*_*d*_. Based on the data presented in Fig 1.b, the decouplexing provides molecularly precise readouts, the AQ model, applying saturating conditions, predicts absolute molar quantities of analytes, and as a consequence, it is noteworthy that the sensitivity of the PICO assay is exclusively constrained by the compartment size (number of compartments), Fig B3 and C8).

### Proof of absolute quantification

To validate the AQ model underlying the absolute quantitative PICO assay, a comprehensive set of experimental models was developed. Fig 1.d presents a both internally and externally verifiable experimental setup termed triangular PICO (3-cplex assay), in which three distinct PICO antibodies concurrently quantify the analyte. Noteworthy, that under saturating conditions, all three antibodies bind to the analyte simultaneously. As a verification, under saturating conditions, all three ABPs must yield equivalent absolute quantities internally. These internal measurements should also align with the concentration of an externally verified recombinant analyte standard. Fig 1.d shows the concept of the AQ verification experiments of rErbB2 (his-HER2) analyte under saturating condition with the verification of saturation (Fig 1.e), the calculated AQ plots of all three ABPs and all dilutions of analyte (Fig 1.f), linearity and dynamic range (Fig 1.g-h), and the derived absolute amounts under varied experimental setups (Fig 1.i). Similar AQ experiments using a recombinant polytag analyte (HIS-thioredoxin-STREP protein) described in C6. Both lines of experiments corroborated the validity of the AQ model, affirming that accurate single-molecular readouts are precisely interpreted through the AQ framework. Results also includes adjustments pertinent to labeling efficacy and other factors see Materials and Methods (and Appendix B).

### Validation of PICO in cellular matrix and against external references

To extend applications beyond the analytical recombinant systems, we aimed to address the demonstration of minimal to negligible matrix effects that otherwise could introduce bias due to different assay contexts. In Fig 2, we present data employing the same set of ErbB2 triangular antibodies that were previously utilized and validated in analytical experiments, now in cellular context. Notably, the readings obtained using the anti-HIS tag antibody need to be consistently negative since the anti-HIS tag antibody lacks epitope on natural ErbB2 (Fig 2.b). These zero readings encompass the no-cell controls, an antibody only negative control (ABC), which expected to be zero, as well. Various dilutions of lysed BT474 and MCF7 cells measured with anti-HIS-tag result in zero couplexes. (Fig 2.c-d-e). The dataset not only demonstrates the absence of the zero background (ABC values) but also validates the presence of minimal to no matrix effects for challenging lysed cellular context (anti-HIS ABPs). Furthermore, according to Fig 2.g in lysed cellular context the spiked-in amount of rErbb2 was recapitulated. We, also, conducted an extensive AQ PICO measurement (n = 178), comparing results against a commercial ELISA, using BT474 and MCF7 cells with two different ErbB2 ABPs, also by different operators (Fig 2.f). Importantly, the absolute cellular amounts of natural ErbB2 measured were consistently in agreement within a given cell type, especially for MCF7, while at extreme high Erbb2 concentration values, in BT474, significantly differed, though with a low fold changes. To prove specificity of PICO regardless of the assay context, we conducted an ErbB2 spike-in experiment (Fig 2.g). In this experiment, quantified recombinant ErbB2 was added to lysed cells. All absolute quanties, including the amount of spiked-in rErbB2, and the measured natural ErbB2 amounts, were found to be accurate and summable, thus confirming the specificity of determining correct absolute amounts in PICO, irrespective of the formidable and chemically diverse environment of lysed cellular contexts.

**Figure 2:**
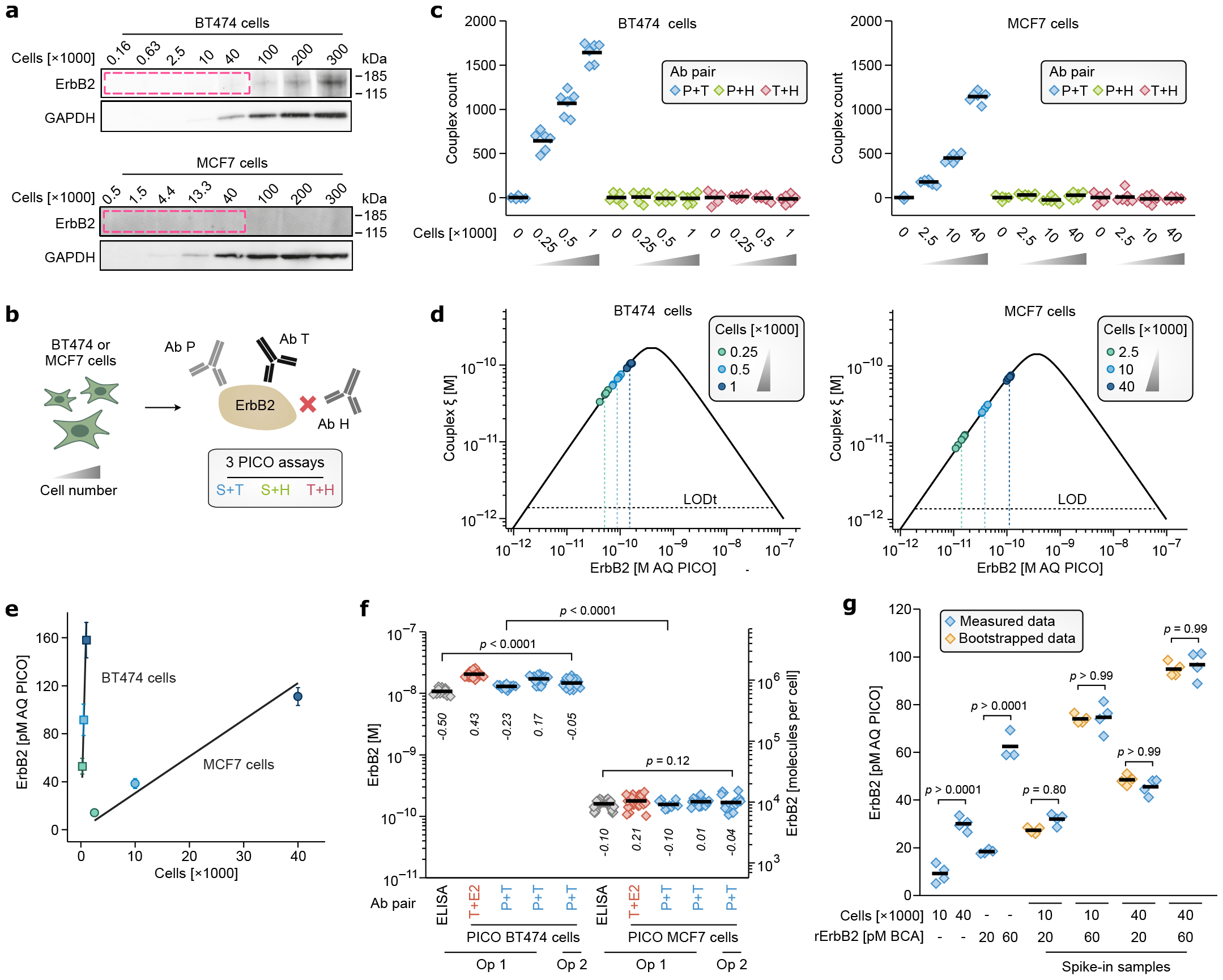
**a**. Western blotting involved loading varying cell quantities of BT474 and MCF7 cells as indicated, followed by immunodetection using trastuzumab (anti-ErbB2) and anti-GAPDH antibodies as loading control (detailed in Material and Methods, Section D.7). For BT474 (ErbB2 over-expressing breast cancer cellline) the ErbB2 signal diminished to an undetectable level at approximately 40,000 cells per lane. Notably, ErbB2 expression in MCF7 cells was scarcely detectable even at 300,000 cells, while the loading controls are comparable. BT474 cells express 1,940,000 ERbB2 proteins per cell (qFACS) [29], while MCF7 exhibits approximately 100-fold lower expression (Western) [30]. Considering a limit of detection (LOD) at 40,000 cells for BT474, MCF7 falls below the LOD of Western at 300,000 cells. **b**. ERbB2 triangular PICO assay with bystander antibody. See also Fig 1.d, note that Ab H was used here as a bystander antibody - BAB, as an internal negative control, as natural ErbB2 has no HIS-tag. **c**. Absolute number of couplexes measured of ErbB2 PICO using different number of lysed BT474 and MCF7 cells (each 6 parallels), as indicated, and ABPs, all assayed at *λ* = 0.15, as described in Material and Methods (D.5). An analyte-free negative control was used, and expected to be zero. Data is normally distributed (Shapiro-Wilk, *p >* 0.1, *n* = 144), the cell measurements of P+T ABP are all significantly different from zero (t-test, *p <* 0.001, *n* = 36), while BAB pairs are not significantly different from zero (t-test, *p >* 0.1, *n* = 108), means are indicated. The bystander antibody (BAB) measuring unspecific binding, the consistently zero BAB signals confirm zero background. **d**. AQ curves of BT474 and MCF7 of ErbB2 PICO based on Fig. data. The Y-axis represents the concentration of couplexes, and the X-axis denotes the absolute molar concentration of the analyte in the binding reaction. The mean concentration of the antibodies of T and P for the BT474 assay is 3.61 × 10^−10^ ± 4.46 × 10^−11^ M (n=46) and for the MCF7 3.25 × 10^−10^ ± 3.56 × 10^−11^ M (n=40), saturating conditions. The calculated absolute concentrations effectively replicate the cell dilution series (data not shown), and yielding estimates of 8.88 × 10^5^ ± 1.50 × 10^5^ ErbB2 per cell for BT474 and 9.86 × 10^3^ ± 3.19 × 10^3^ ErbB2 per cell for MCF7, corresponding a 90.1× of relative expression difference, confirming previous findings [30]. The slight departure of BT474 expression form qFACS data (∼ 0.45×) is explained by biological variability or reproducibility problems (for qFACS [31]), also PICO is highly consistent between experiments, and with ELISA, see also (Fig. 2.f). **e**. Absolute concentration vs. cell number of data Fig.**c**. The LOD calculation was conducted according to Wenzl et al.[28],and a LOD of 17.31 ± 15.96 BT474 cells and 982.53 ± 856.29 MCF7 cells were determined, respectively. Similarly, LOQs were 75.14 ± 52.6 BT474 cells and 4264.18 ± 2825.76 MCF7 cells. **f**. The reproducibility of PICO. ErbB2 proteins of BT474 and MCF7 cells were measured with PICO, including varied of ABPs and operators and verified against ELISA. The double Y-axis represent ErbB2 concentrations (in the binding reaction) and proteins per cell. Log2 fold differences were calculated as (sample mean) / (group mean), and also indicated. All (PICO) average copy numbers per cell of ErbB2 were 1.02 × 10^6^ (±2.10 × 10^5^), and 1.06 × 10^4^ (±2.96 × 10^3^) (*n* = 178), for BT474 and MCF7, respectively. BT474 cells expressed approximately 96.52 (±25.32) times more ErbB2 than MCF7 cells, a significant difference (F(9,220)=941.64, p-value<0.0001, n=229). ANOVA revealed no statistical difference between MCF7 samples (F(4, 135)=1.86, p-value = 0.121, n=139), while BT474 cells exhibited significant differences (F(4,85)=53.03, p-value < 0.0001, n=89), albeit with minimal deviation (*CV* = 20.56%) and low fold differences, attributable to biological variations. ELISA measured an ErbB2 expression per cell 6.49 × 10^5^ ± 8.34 × 10^4^, 0.69 times less than average PICO data, *p* = 0.0026, *n* = 66, and 9.45 × 10^3^ ± 1.57 × 10^3^, 1.06 times less then PICO, *p* = 0.19, *n* = 118, for BT474 and MCF7, respectively. The individual molar ErbB2 of each group from left to right are 10.8 (±1.38) nM, 20.4 (±2.29) nM, 13.0 (±1.01) nM, 17.1 (±2.37) nM, 14.7 (±2.25) nM, 157 (±26.0) pM, 194 (±56.3) pM, 157 (±21.5) pM, 169 (±40.3) pM, 164 (±52.9) pM. Altogether PICO showed high precision and reproducibility, especially for MCF7 cells, BT474 has higher ErbB2 expression per cell and consequently more effected with sampling errors of cell counting. **g**. The specificity of PICO. An absolute quantitative spike-in experiment was carried using two dilutions of MCF7 cells (10,000 and 40,000 cells, representing the theoretical concentrations of 39.3, and 157.5 pM, respectively) and rErbB2 protein (60 and 20 pM). Samples were as indicated (*n* = 4), and were compared to bootstrapped values. ANOVA were applied to compare bootstrapped combinations of MCF7 and rErbB2 concentrations demonstrating no significant differences (*p <* 0.05) in all comparisons. The spike-in experiment demonstrates robust specificity, allowing unbiased measurement of rErbB2 levels in the presence of endogenous ErbB2 and lysed cell material, maintaining concentration additivity.

PICO was compared to SIMOA [11], Fig C8, and Western blot (Fig 2.a, 3.a). SIMOA is showing comparable molar sensitivities (LODs are fM for SIMOA and 10s of fM for PICO, Fig C8.b) under conditions applying comparable number of beads and compartments, respectively, however, comparing them directly is challenging because of inherent differences. PICO uses a sample volume 15 times smaller, but if compensated for this, they exhibit equal sensitivity (≈2 fM). Additionally, PICO has a sensitivity-limiting dilution step before detection. Commercial dPCR instruments used for PICO have less compartments than the applied beads in SIMOA, which is a practical limit for sensitivity of PICO rendering it in the 100s of fM - pM range, however with hyperwelling (merging physical dPCR wells) (Fig C8) this boundary is permeable. Also, high-compartment-sized dPCR technologies are already on the horizon [12], offering theoretical attomolar sensitivities. Unlike PICO, SIMOA does not function as a single-molecule counting assay, as verified and evident from the absolute amounts detected (see LOD [molecules] in Fig C8.c), and it is not background free [11]. Fig 2.e shows the cellular LOD of ERbb2 PICO (17 ± 16 cells for BT474, 1.6 × 10^6^ ErbB2 copies per cell, and 983 ± 856 cells for MCF7, 1.2 × 10^4^ ErbB2 copies per cell) against Western blotting, Fig 2.a, (LOD of 40,000 cells for BT474) clearly demonstrating an outperformance by a factor of more than 2000 times. PICO, a liquid phase assay, has no lower assay volume limit, allowing for adjustment of binding conditions through volume downscaling. This reduction in volume decreases antibody quantities at the same concentration, enabling a significant reduction in the extent of dilution, and pushing the assay’s absolute sensitivity from the attomole regime down to the low zeptomole level, allowing detection of just a few thousand or even below of molecules. This adaptability allows PICO to achieve sensitivities down to scope a single-cell while maintaining it lossless absolute quantitative performance (unpublished data).

### Multiplex PICO measuring cellular stoichiometry with high precision

Based on the proven absolute quantitative PICO, a novel 8-plex PICO assay was designed, which aimed to partly replicate previously reported complex pathway studies [25], and including many proteoforms, extending the PICO assay toward highly parallel regimes. Fig 3 presents an 8-plex, split-readout dPCR-based PICO assay, with the split readout of S6K1, p4EBP1 and ErbB2, ErbB3, ErbB2:ErbB3 interaction, respectively (Fig 3.d) including protein, phosphoprotein and interaction targets. The assay detects 6 targets (the ErbB2:ErbB3 interaction measured with two different ABPs), and 6 BAB pairs (bystander antibody controls) i.e. confirmatory to zero readouts (Fig 3.e-f). Combinatorial multiplexing (cplex) exerts exceptional control over PICO reactions. The 8-plex split-readout PICO, taking into account the cell dilutions as distinct samples, includes 18 non-target BAB readings, and 12 no-cell negative controls, which are essential for leveraging dPCR clustering biases (see Methods). Furthermore, there are 18 target readouts. As the purposely applied pertuzumab cannot bind to ErbB2:ErbB3 complexes [23] providing a unique opportunity to confirm the zero PPI readings by comparing them to trastuzumab ABPs, both of them even based on two different ABPs readouts (Fig 3.f). Furthermore, the two trastuzumab-based readouts are non-significatly different (Fig 3.i) along all conditions. Nevertheless, the functionality of pertuzumab is intact per ErbB2 readouts (Fig 3.g-h). Small, down to 1.33-fold, and large fold changes are precisely measurable between treatments (Fig 3.g-h). The absolute quantitative results enable unrestricted comparisons revealing the quantitative stoichiometry of protein complexes and phosphoproteins, even between cells, especially for p4EBP1 (Fig 3.c) and ErbB proteins (Fig 3.g-h), elegantly proposing the rate-limiting nature of ErbB3 expression in the ErbB2:ErbB3 complex formation in MCF7 cells. Regarding dactolisib treatment of MCF7 (low ErbB2) and BT474 (high ErbB2) cells, 8-plex PICO recapitulates earlier [25] findings of increased ErbB2 and ErbB3 expression, however, while ErbB2:ErbB3 is confirmatively increased in MCF7 cells as well, in BT474 it is reduced to an undetectable level, falling below the limit of detection (LOD) of 181 molecules per cell. In contrast, untreated BT474 cells exhibited detectable ErbB2:ErbB3 interaction.

**Figure 3:**
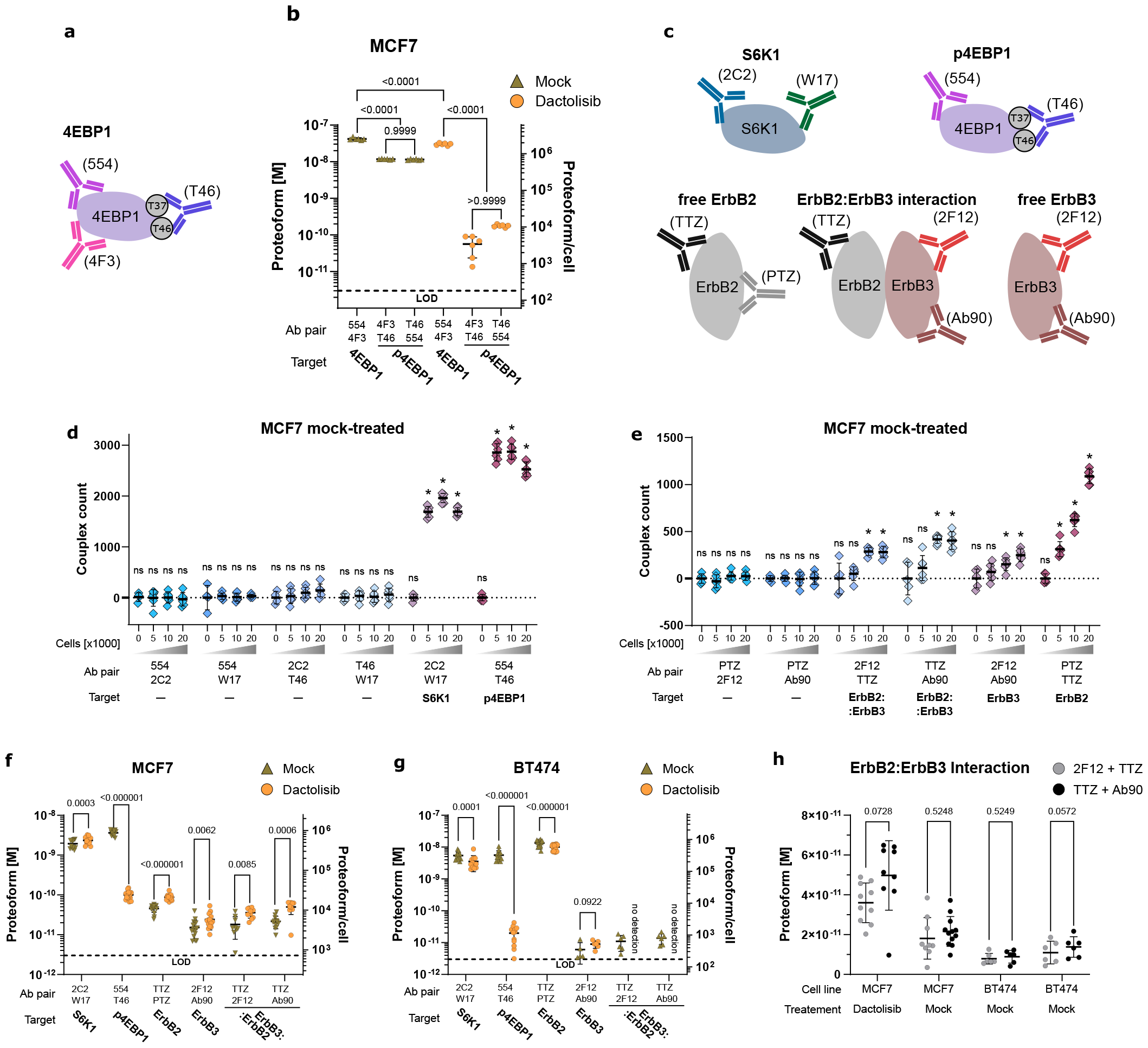
8-plex PICO assay detecting different proteoforms. **a**. Triangular detection of 4EBP1 and its phosphorylation. Note that the redundant absolute quantification of 4EBP1 phosphorylation (measured by two ABP). **b**. Detection of 4EBP1 protein and its T37-T46 phosphorylation using mock and dactolisib-treated MCF7 cells. After treatment 40k MCF7 cells were lysed and PICO absolute quantified. The y-axis is in molar concentration, while the right axis is proteoforms per cell, the LOD indicated. The mean number of proteoforms per cell (ppc), from left to right, are 2530000 ± 220000 ppc, 707000 ± 21000 ppc, 694000 ± 31000 ppc, 1810000.00 ± 90000 ppc, 3400 ± 2200 ppc, 10700 ± 700 ppc. Based on ANOVA (F(5, 30) = 643.0, p < 0.0001, n = 36), dactolisib significantly perturbs both protein and phosphorylation levels (both p < 0.0001, n = 12) the latter showing a significant difference in phosphorylation (27% versus 0.4%). 4EBP1 phosphorylation as detected by the two different ABPs showed no significant difference between mock (p = 0.9999, n = 12) and dactolisib-treated (p >0.9999, n = 12) MCF7 cells demonstrating concordant absolute quantitative results. **c**. Depictions of PICO assay used for the detection of the cytosolic protein S6K1 and p4EBP1 along with the detection of ErbB2, ErbB3, and ErbB2:ErbB3 interaction. The 8 antibodies were concurrently applied (8-plex PICO), however, detected by two different amplification primer pairs in separate dPCR reactions, S6K1, 4EBP1-phosphorylation and ErbB2, ErbB3, respectively. Note the inability of pertuzumab to bind to the interacting protein complex. **d,e**. Couplex counts per dPCR reactions of 8-plex PICO using 0, 5000, 10000, or 20000 lysed MCF7 cells (each 6 replicates), ABPs are indicated, normalised and *λ*-corrected values (see methods). All antibodies were applied in parallel see (d), (e) S6K1 and p4EBP1, ErbB2 and (f) ErbB2, ErbB3 and ErbB2:ErbB3 interaction. The asterisk (*) indicates a p-value of less than 0.05. No sign indicates nonsigficant difference from zero ((e) - normality test Shapiro-Wilk test p = 0.18839, n = 91, two-sided one-sample t-test t(90) = 1.62542, p = 0.10757, n = 91; (f) - normality test Shapiro-Wilk test p = 0.11861, n = 66, two-sided one-sample t(65) = 0.21725, p = 0.82869, n = 66). **f**. Mock or dactolisib-treated MCF7 cells (20k cells, low ErbB2 expression) were PICO absolute qualified using the 8-plex PICO assay described above (d), all comparisons are in mock-dactolisb order. S6K1 levels were marginally, but significantly different between mock and dactolisib treatments (468,000 ± 112,000 ppc and 567,000 ± 128,000 ppc, respectively) indicating high precision, but presumably no or minimal biological significance (t(11) = 4.845, q = 0.000347, n = 24). The pronounced and expected effects of dactolisib treatment on p4EBP1 were confirmed (875,000 ± 130,000 ppc and 24,200 ± 5,700 ppc, respectively, 36-fold difference, t(10) = 20.54, q < 0.000001, n = 29). Note that the level of p4EBP1 is susceptible to variations in culturing conditions. There was an expected, known difference in ErbB3 level, confirming literature (see main text) having 3,670 ± 1,980ppc and 5,810 ± 2,640 ppc, respectively, a 1.6-fold increase, t(13) = 2.789, q = 0.006206, n = 32. ErbB2 levels were also significantly different (10,940 ± 2,160 ppc and 21,100 ± 3,400 ppc, respectively, an 1.9-fold increase, t(17) = 11.3, q < 0.000001, n = 36). ErbB2:ErbB3 interaction was significant between treatments, however, consistent between the two antibody pairs TTZ-2F12 and TTZ-Ab90, respectively, having 4,370 ± 2,520 ppc and 8,670 ± 2,400 ppc, a 2-fold increase, t(6) = 2.959, q = 0.008528, n = 19 and 5,200 ± 1,810 ppc and 12,000 ± 4,200 ppc, an 2.3-fold increase, t(8) = 4.848, q = 0.000644, n = 21. In mock-treated MCF7 cells, approximately 44 **g**. Mock or dactolisib-treated BT474 cells (40k cells, high ErbB2 expression) were PICO absolute qualified using the 8-plex PICO assay described above (d), all comparisons are in mock-dactolisb order. S6K1 levels were different, having 326,000 ± 95300 and 193,000 ± 60,700 ppc, t(11) = 5.91, q = 0.000034, n = 24, confirming precision, but marginal changes. p4EBP1 levels were consistent with the dactolisib treatment, 337,000 ± 125,000 and 1,190 ± 716 ppc, t(11) = 9.32, q <0.000001, n = 24 exhibiting a high sensitivity to dactolisb and shows 283-fold phosphorylation difference. ErbB2 levels, contrary to MCF7 data, show a decrease of ErbB2 level upon dactolisib treatment, 808,000 ± 177,000 and 604,000 ± 128,000 ppc, a 1.33-fold decrease, t(17) = 9.20, q < 0.000001, n = 36. While ErbB3 protein levels were highly concordant with MCF7, having 312 ± 244 and 541 ± 142 ppc, an 1.7-fold increase, (t(4) = 2.20, q = 0.023303, n = 12). Nevertheless, ErbB2:ErbB3 interaction was only detectable in mock-treated cells, having concordant, non-significantly (see below) different levels with the TTZ-2F12 and TTZ-Ab90 ABPs and was 662 ± 343 and 831 ± 311 ppc, respectively, the LOD was 200 ppc. In dactolisib-treated BT474 cells, the ErbB2:ErbB3 interaction is extremely low or absent, as evident from the data. While, in mock-treated cells, approximately 0.09% of ErbB2 interacts with ErbB3, while nearly 100% of ErbB3 engages in interaction with ErbB2, replicating its rate-limiting behaviour seen in MCF7 cells. **h**. Highly concordant results between TTZ-2F12 and TTZ-Ab90 ABPs detecting the ErbB2:ErbB3 interaction from left to right are t(7) = 2.588, q = 0.0728, n = 19; t(8) = 0.8282, q = 0.524873, n = 21; t(5) = 0.6921, q = 0.524873, n = 12; t(5) = 3.690, q = 0.05715, n = 12, demonstrating the self-confirming applicability of such parallel measurement.

## Discussion

Why do we need a new type of immunoassay (IA) among the, to say the least, many [13-15]? The ever-more-capable IAs have achieved very high sensitivities, in both molar, SIMOA and PEA [11, 16], and absolute terms [17], and reached high parallelism OLink [18] and SOMAmers [19]), however, one overlooked feature is still missing — a truly reference-free IA for the absolute, and universally comparable, quantification of analyte. Besides evident practical reasons, absolute quantification of proteoforms is a must for highly parallel cellular stoichiometry [20] waiting to set out to understand intricate cellular workings and exploring maybe just stoichiometrically different views with consequent functional differences [21]; recently, structural studies have opened up views on a vastly diverse proteomic and interatomic landscape and, naturally, on the intriguing stoichiometrics of these diversities [22]. In this context, PICO introduces new paradigms, comprising a compartmentalized, homogeneous, single-molecule assay using a new lossless signal generation principle, which robustly achieves a limit of detection (LOD) of a few or practically zero molecules per reaction, limited only by unavoidable Poisson-based stochasticity. Through its unique physical assay principle of partitioning, read out by decoupexling, and its inherently separate binding and detection steps, PICO stands alone in achieving zero background. The absolute signal is mathematically described introducing practically attainable theoretical chemistry for IAs. The analysis of PICO chemical foundations fosters the dissociation constant agnostic PICO applications using antibodies or any binding agents. Validation experiments, including internal and external verifications, confirm the reliability of absolute quantitative PICO across various experimental setups, including cellular contexts, plus demonstrating never-before-seen, formerly unimaginable features, like combinatorial multiplexing (cplex) exemplified here by triangular, quadratic PICO assays Fig 1.d and 8-plex split Fig C6.e. These higher-order cplex assays facilitates complex readout of proteoforms e.g. complex protein interactions and concurrently the involved interacting proteins are measured (for six ABPs, Fig 3), enabling self-confirming multiple measurements on the very same target molecule, and a vast array of negative (expecting zero signal) control reactions (using BABs or just between non cognate antibodies) exerting strict control over the entire multiplex reaction network. Even structural studies, i.e. the therapeutic antibody of ErbB2, pertuzumab, known having epitope on the interaction surface [23] rendering the ErbB2:ErbB3 interaction signal to be zero, while trastuzumab detects the interaction absolute quantitatively. Compared to existing techniques like SIMOA [24] and Western blotting, PICO exhibits superior sensitivity, especially evident in cellular LODs, outperforming Western’s sensitivity. dSIMOA claims absolute quantification, due to lack of access to the technology it was not possible to demonstrate, nevertheless based on the published data [11] the LOD of dSIMOA is still a few thousand molecules at best and it is inferior to PICO’s capabilities. Upon dactolisib treatment, the increase of ErbB2:ErbB3 interaction was recapitulated [25] in the breast cancer cell line MCF7 using 8-plex detecting down to 181 proteins in cells Fig 3 demonstrating multiplexity, combinatorial multiplexing, the detection of an array of proteoforms, self-confirmation, structural hindering and strict control of specificity. Nevertheless, PICO is homogeneous and solid-phase-free unlike ELISA-based concepts (i.e. [17], [11] and [19]), allowing the adaptability of the assay to ultra-low volumes and allowing PICO to achieve sensitivities in absolute terms down to the scope of a single cell, i.e. detection of zeptomoles were demonstrated in single-cells, while maintaining its absolute quantitative, lossless performance (using a 12 nL PICO reactions, CSHL Single Cell Analyses, Nov 8 - 11, 2023, manuscript in preparation). While others fail to show negligible background, which in turn hampering quantification at the single cell level [26]. The data also demonstrate PICO endurance against matrix effects and exhibiting high specificity Fig 2c. and g., which was further underlined by absolute detection of rErbB2 in spiked-in whole blood (unpublished) or demonstrating zero signal indicating mutually exclusive binding (see above Fig 3 and using anti-p4EBP1 and anti-4EBP1 Abs targeting the same epitope, unpublished). In conclusion, PICO has enormous potential, offering versatile, standardized, precise measurements of proteins, providing insights into the physiologically relevant stoichiometry of disease-related cellular processes, even revealing structural intricacies and hence facilitating the development of therapeutic agents [27].

## Appendix A The rationale behind molecular counting of proteoforms

We introduce a novel immunoassay detection principle known as ‘compartment decouplexing.’ This model holds significant implications for immunoassay applications and has recently been applied to replace co-immunoprecipitation (coIP) in viral receptor research[6]. It serves as the foundation for the establishment of protein interaction coupling (PICO), previously referred to as ‘emulsion coupling’. PICO employs a bi-component approach as utilizing two specific binding antibodies per analyte, defining the detection unit as the ternary complex formed involving the antibodies and the target antigen, referred to as a ‘couplex’. The detection concept capitalizes on the differential distribution of antibodies in the compartments, which arises from the differential distribution of the free antibodies versus antibodies bound to couplexes. Maintaining the average number of antibodies per compartment, generally less than 1, denoted as *λ*, the distribution of antibodies remains mathematically tractable, as it is always possible to determine the number of couplexes in the compartments regardless of the number of antibodies present. During the readout, compartments are stratified based on antibody content, they might contain one type of antibody, the other, or both. Importantly, a compartment containing two antibodies does not always indicate the presence of a couplex due to potential random collocation. Compartments with both antibody types are labeled as duplex compartments, altogether leading to three distinct output metrics - two single types and the duplex type. Simultaneously, the total compartment count is also determined. Our model posits that these four metrics, more specifically their compartment negative counterparts see below, as **X** = (*X*_*L*−_, *X*_*R*−_, *X*_−−_) and *m* - number of compartments -, when considered collectively, offer a robust probabilistic framework to ascertain the number of couplexes within the compartments.

### A.1 The model of compartment decouplexing

There are *n*_*L*_ molecules of antibody L (left) - the term left and right used to differentiate the single types of antibodies, these terms does not imply differences between the antibodies, and *n*_*R*_ molecules of antibody R (right) and *n*_*C*_ couplexes are formed from these antibodies, and as stated above, each couplex contains one antibody L, and one antibody R molecule. As a simple consequence, *n*_*uL*_ = *n*_*L*_ − *n*_*C*_ the unbound antibody L (*uL*), and *n*_*uR*_ = *n*_*R*_ − *n*_*C*_ the unbound antibody R (*uR*), while *n*_*C*_ is the couplex (*C*) molecules. The *n* = *n*_*uL*_ + *n*_*uR*_ + *n*_*C*_ = *n*_*L*_ + *n*_*R*_ − *n*_*C*_ molecules are distributed in *m* compartments (*m* → ∞), such that each antibody molecule, independently of each other, has equal probability 1*/m* to fall in each compartment. The number of compartments *m* is known. Let

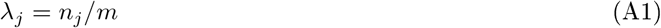

be the average number of type *j* molecule per compartment (*j* = *uL, uR, C*). For the sake of convenience let us introduce the ordered index set N = {*uL, uR, C*}. In the sequel we consider the limiting case when *m* and *n*_*j*_ (*j* ∈𝒩) tend to infinity in such a way that *λ*_*j*_ tends to a finite positive limit *κ*_*j*_. A compartment is positive for L (respectively, for R), if it contains either *uL* or *C* (respectively, either *uR* or *C*). A compartment is positive for both L and R, if it contains *C*, or both *uL* and *uR*. The model parameters to be estimated are *ϑ* = (*n*_*uL*_, *n*_*uR*_, *n*_*C*_), or equivalently, *ϑ* = (*n*_*L*_, *n*_*R*_, *n*_*C*_).

For simplicity, the number of compartments, that are negative for the respective measures of antibodies listed above, are used, thus the vector of observed data is (*X*_*L*−_, *X*_*R*−_, *X*_*LR*−_), where *X*_*L*−_ is the number of compartments negative for antibody L, i.e. they contain neither *uL*, nor *C*. Similarly, *X*_*R*−_ is the number of compartments negative for antibody R, i.e. they contain neither *uR*, nor *C*. Finally, *X*_*LR*−_ is the number of compartments negative for at least one antibody, i.e. they do not contain *C*, furthermore at least one of *uL* and *uR* are missing. Conveniently introducing *X*_−−_ = *X*_*L*−_ + *X*_*R*−_ − *X*_*LR*−_ for the number of compartments negative for both antibodies, i.e. they are completely empty, the random vector of observed data is **X** = (*X*_*L*−_, *X*_*R*−_, *X*_−−_). To refer to the coordinates of this vector in a simple way, we introduce the ordered index set ℳ= {*L*−, *R*−, −−}.

To estimate the parameter vector *ϑ* from the observed data **X** the method of moments was used, i.e for the realization **x** of the data we solve the system of equations:

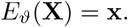

We remark that the measurement could be repeated several times, with the same composition of molecules. In this case, the right-hand side of the above equation is replaced by the average of the measured data.

All three coordinates of **X** count the number of compartments not containing the respective antibody molecules. Such random variables are well-studied in the literature, see e.g. the details in Chapter 1 of [32]. Define *µ*_*i*_ = 𝔼 _*ϑ*_ (*X*_*i*_)*/m* = 𝔼(*X*_*i*_)*/m, i* ∈ ℳ. Note that in most formulae, we will suppress the dependence on the parameter *ϑ*. The expected value of *X*_*L*−_*/m* is

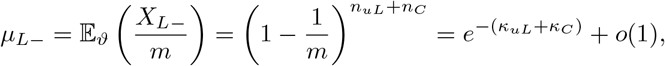

and similarly, for the other two coordinates of **X**:

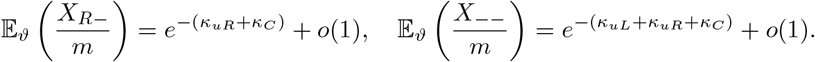

Thus, an approximate moment estimator for 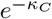 is

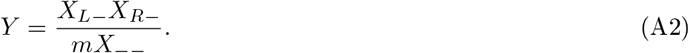

From equation A2, and introducing *ξ* as the number of couplexes, −*m* ln (*Y*) gives an estimator for the number of couplexes detected. For experimental confirmation of this derivation of *ξ* a simulation experiment was carried at varied *λ*_*uL*_ = *λ*_*left*_, *λ*_*uR*_ = *λ*_*right*_ and *ξ*, see in Fig A1 (details in caption). Next step is to prove the normality of *ξ*.

**Figure A1:**
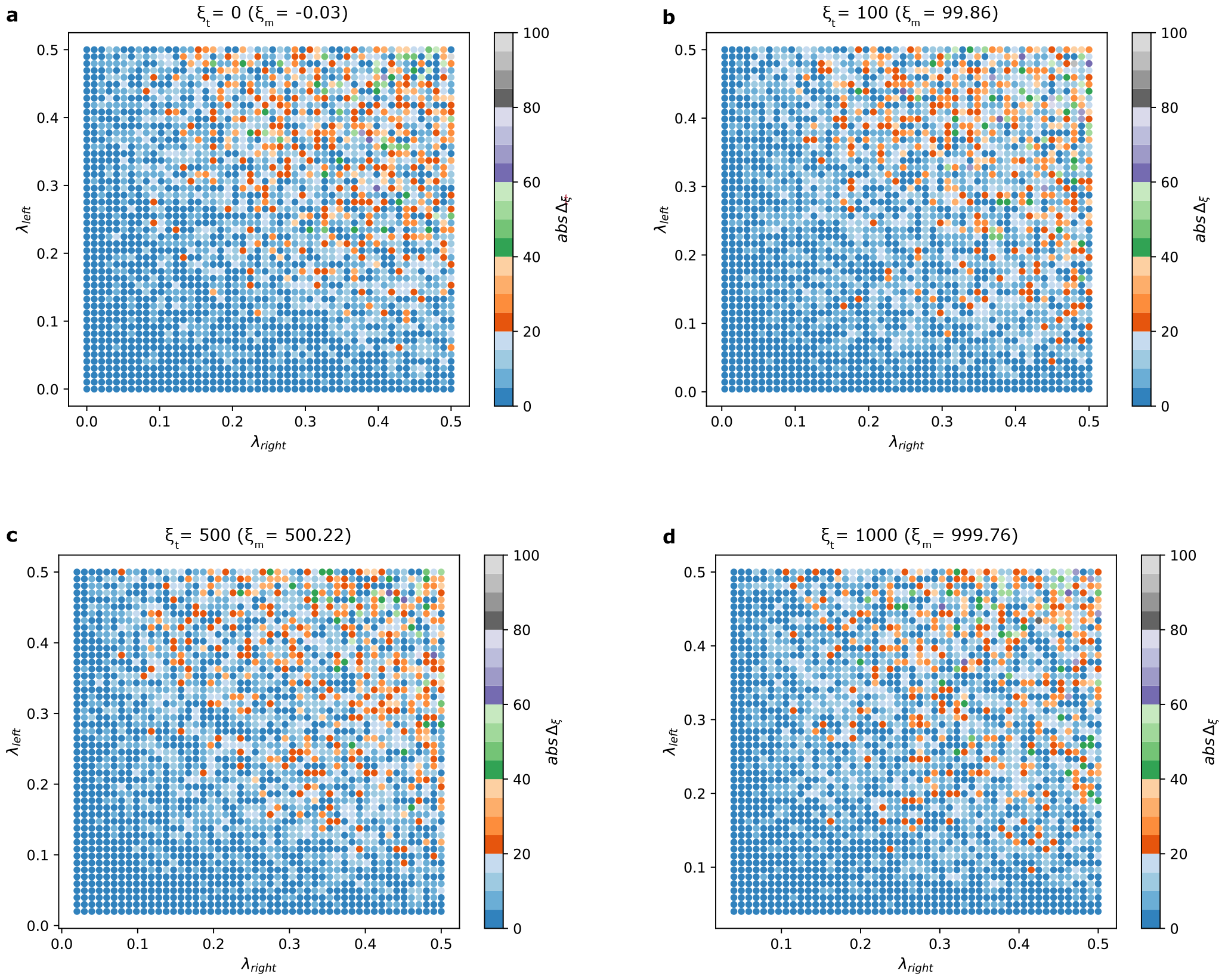
Numerical modeling of decouplexing. To simulate decouplexing a sum of three independent Poisson processes was applied using 25,000 partitions. Two were used to generate Poisson distributions varied according to parameters *λ*_left_ and *λ*_right_ representing the two free antibodies, both incremented over a range of *λ* of [0, 0.5] in 50 equal steps. The third process pertained to the parameter *ξ* as number of couplexes assigning values of 0, 100, 500, and 1000 (theoretical values of *ξ*), see subfigures from A through D, respectively. The aggregation of these processes conforms to a Poisson distribution (proof not shown). The X and Y axes delineate the *λ* values. The number of couplexes is estimated using Eq. *A*2, over 10 simulations for each grid point and the delta (*abs*Δ_*ξ*_) was calculated as the absolute difference of the mean and the theoretical *ξ* value and indicated as a color gradient. Above each subplot the overall mean of *ξ* of all simulations (*n* = 25000) is shown. The results unambiguously show the agreement between of the simulated and theoretical *ξ* values, regardless of the value of *λ* or the theoretical *ξ*, and confirms the validity of the model. Although the *abs*Δ_*ξ*_ is influenced by the *λ*, it consistently remains below 20 couplexes when *λ* ≤ 0.2, similarly it largely remains below 40 when *λ* ≤ 0.5. It is noteworthy that the distribution of absolute deviation remains largely unaffected by *ξ* across the Subfigures A to D. This trend underscores the inference that a chosen *λ* retains its constant absolute deviation universally, irrespective of the *ξ* value. Consequently, for a fixed *λ* pair, the coefficient of variation decreases as *ξ* increases. The variance has a defined mathematical form. see Eq. A3.

### A.2 Normality

Let us consider the asymptotic distribution of **X**. To simplify the formulae, introduce the function

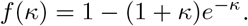

#### Theorem 1

*uppose m and n*_*j*_ (*j* ∈ 𝒩) *tend to infinity in such a way that λ*_*j*_ *tends to a finite positive limit κ*_*j*_. *Then the joint distribution of the random vector*

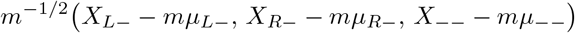

*converges to a* 3*-variate normal distribution with zero mean and variance*

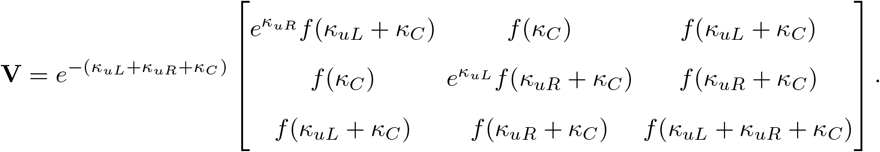

*Proof*. For *k* = 1, …, *m* and *j* ∈ 𝒩 introduce *ξ*_*k,j*_ as the number of type *j* molecules in compartment *k*. Then

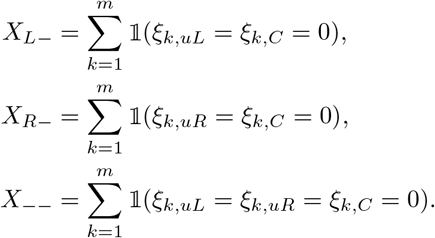

Let *Z*_*k,j*_, 1 ≤ *k* ≤ *m, j* ∈ 𝒩, be independent Poisson distributed random variables, *Z*_*k,j*_ with parameter (i.e., mean) *λ*_*j*_. Moreover, let

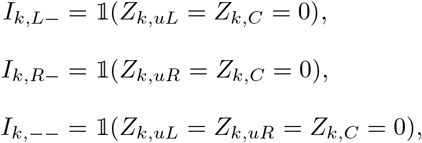

and 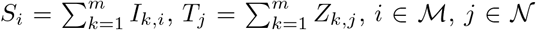. Then the joint distribution of *ξ*_k,j_, 1 ≤ *k* ≤ *m*, j ∈ 𝒩, coincides with the conditional joint distribution of *Z*_*k,j*_ supposed that *T*_*j*_ = *n*_*j*_, *j* ∈ 𝒩 . Hence, the joint distribution of *X*_*i*_, *i* ∈ M, coincides with the conditional joint distribution of *S*_*i*_ supposing that *T*_*j*_ = *n*_*j*_, *j* ∈ 𝒩 . Thus we are interested in the joint asymptotic distribution of the random variables *S*_*i*_, *i* ∈ ℳ, *T*_*j*_, *j* ∈ 𝒩.

Consider the i.i.d. random vectors

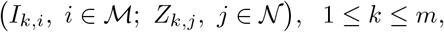

with integer coordinates. It is easy to see that they have mean

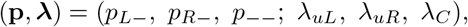

where 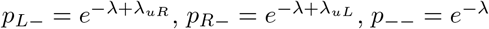, *λ* = *λ*_*uL*_ + *λ*_*uR*_ + *λ*_*C*_ = *n/m*, and variance

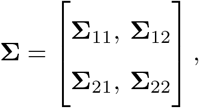

where

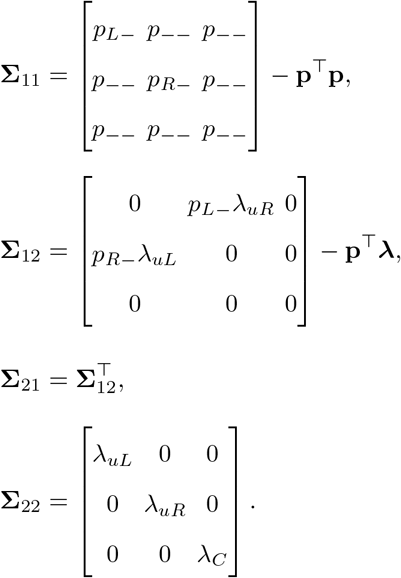

Indeed, 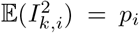, and 𝔼(*I*_*k,i*_*I*_*k,j*_) = *p*_−−_ if *i* ≠ *j* (*i, j* ∈ ℳ), moreover 𝔼(*I*_*k,L*−_*Z*_*k,uR*_) = *p*_*L*−_*λ*_*uR*_, 𝔼(*I*_*k,R*−_*Z*_*k,uL*_) = *p*_*R*−_*λ*_*uL*_, and 𝔼(*I*_*k,i*_*Z*_*k,j*_) = 0 in all remaining cases (*i* ∈ ℳ, *j* ∈ 𝒩).

Clearly, if *m* and *n*_*uL*_, *n*_*uR*_, *n*_*C*_ tend to infinity in such a way that lim *λ*_*j*_ = *κ*_*j*_, then the variance above conveges to a 6 × 6 matrix **Δ**, which can be defined similarly, but with *λ*_*j*_ replaced by *κ*_*j*_ everywhere.

By the multivariate central limit theorem the limit distribution of

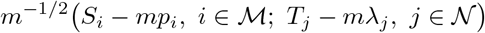

is sixvariate normal, with mean zero and variance **Δ**. Let *η* : ℝ ^ℳ^ × ℝ ^𝒩^ → ℝ denote the corresponding density function. Then Theorem 3 of [33] says that

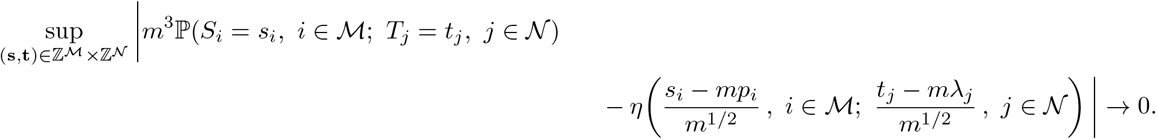

In fact, Davis’ theorem would require the existence of a 6-dimensional integer vector (**s, t**) such that 𝕡 (*I*_*k,i*_ = *s*_*i*_, *i* ∈ ℳ; *Z*_*k,j*_ = *t*_*j*_, *j* ∈ 𝒩) *>* 0, and this probability should remain positive if exactly one of the six integers *s*_*i*_, *t*_*j*_ is increased by 1. This condition aims at excluding the case where the distribution of (*S*_*L*−_, *S*_*R*−_, *S*_−−_, *T*_*uL*_, *T*_*uR*_, *T*_*C*_) is concentrated on a sparse lattice. Here this condition isn’t met, but the remark after Theorem 2 of [33] shows that it is still sufficient if this condition holds for a (fixed size) sum of random vectors. Now, the sum of as few as four summands already possesses the required property with (**s, t**) = (1, 1, 0, 3, 3, 3). (indeed,

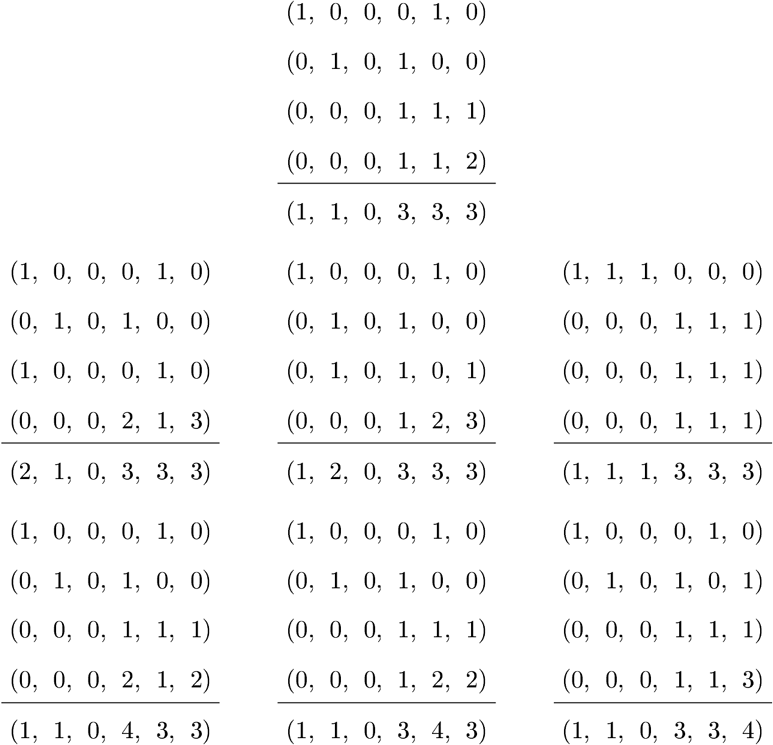

In each table, the four rows above the horizontal line are vectors that can be taken by (*I*_*k,i*_, *i* ∈ ℳ; *Z*_*k,j*_, *j* ∈ 𝒩) with positive probability. The sum of the four row vectors can be found below the horizontal line.)

On the other hand, since *T*_*uL*_, *T*_*uR*_, *T*_*C*_ are independent and their distributions are Poisson with parameter *n*_*uL*_, *n*_*uR*_, *n*_*C*_, resp., it immediately follows that

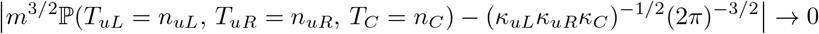

(the term on the right-hand side of the minus sign is just *φ*(**0**) where *φ* is the density function of the threevariate normal distribution with mean zero and variance **Δ**_22_).

Consequently,

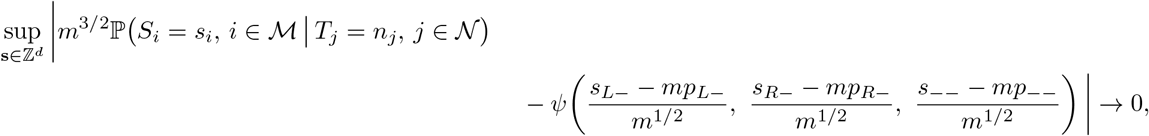

where *ψ* : ℝ ^ℳ^ → ℝ is the density function of the threevariate normal distribution with mean zero and variance 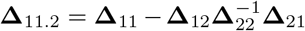 (this is the conditional distribution of ***ζ***_1_ | ***ζ***_2_ = **0** where (***ζ***_1_, ***ζ***_2_) ∼ Normal(**0, Δ**)).

Thus we obtain the following local limit theorem.

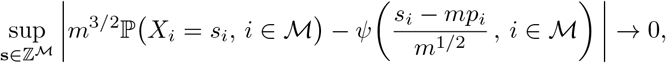

Note that **Δ**_11.2_ = **V**. Now the corresponding global limit theorem, which was to be proved, follows by standard arguments. □

Note that from the proof of Theorem 1 one can easily see that

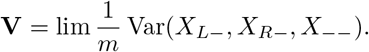

Thus the asymptotic normality of (*X*_*L*−_, *X*_*R*−_, *X*_−−_) is just what one expects. They are sums of identically distributed random vectors, where the summands are not independent but exchangeable. The limit theory for infinite sequences of exchangeable random variables is quite well explored, because they are conditionally i.i.d. by de Finetti’s theorem. This is no more true for finitely exchangeable collections; this is why our result is not so obvious, though the limit distribution is no surprise.

Finally, the following well-known lemma will lead us to the asymptotic normality of *Y* .

#### Lemma 1

*Let* (***ζ***_*m*_) *be a sequence of d dimensional random vectors*, (**a**_*m*_) *a sequence of d dimensional vectors converging to* **a** ∈ ℝ^*d*^ *as m* → ∞, (*b*_*m*_) *a sequence of postive real numbers tending to infinity, and suppose that b*_*m*_(***ζ***_*m*_ − **a**_*m*_) *converges in distribution to* Normal(**0, V**). *Let g* : ℝ^*d*^ → ℝ^*k*^ *be continuously di erentiable in a neighborhood of* **a**. *Then*

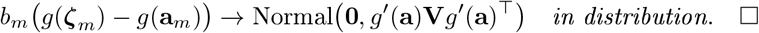

To apply the lemma, choose *ζ*_*m*_ = **X***/m*, then

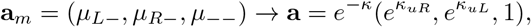

where *κ* = *κ*_*uL*_ + *κ*_*uR*_ + *κ*_*C*_, moreover *b*_*m*_ = *m*^1*/*2^. Then the convergence in the Lemma is just what was proved in Theorem 1. The function *g*(*x, y, z*) = *xy/z* satisfies the conditions of the Lemma, therefore

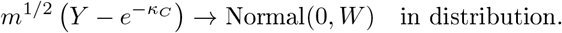

By Lemma 1, the estimator for the number of couplexes detected, i.e. −*m* ln(*Y*), is also approximately normally distributed with variance 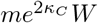.

### A.3 Variance

where the asymptotic variance is calculated to be

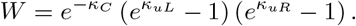

When we want to estimate *W*, we plug in the estimates of *κ*_*j*_, and we get

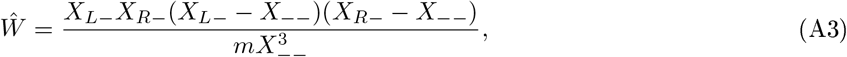

## Appendix B Rationale for absolute quantification of proteoforms

Absolute quantification facilitates the accurate assessment of molecular concentrations, aiding in the comprehensive examination of biological systems, including expression levels, quantitative dynamics of the proteome, protein-complex stoichiometry, degradation rates, post-translational modifications (PTMs), and even network-level quantitative changes shedding light on their material flow and information processing capabilities. Absolute quantification also plays a crucial role in the evaluation of biological models and holds potential clinical applications in disease diagnosis and therapeutic development [27]. The diverse array of proteoforms poses challenges for reference-dependent methods, often requiring the difficult-to-produce quantitative standards, particularly in cases involving protein interactions or modifications. Nonetheless, the complexity of biological systems presents several challenges. Proteins exhibit significant variations in their physicochemical properties, necessitating the design of chemically non-selective assays, and having vastly different cellular abundances, often differing by orders of magnitude. Additionally, proteins are situated within intricate, organized, and compartmentalized environments, making their assessment a challenge, especially membrane proteins or cell organelles. Due to these hurdles, reference-free absolute quantitative analytical methods are scarce in the study of biological systems, and mainly are based on mass spectrometry techniques. Inductively coupled plasma (ICP) ionization is a uniquely robust, reference-free quantitative method and enables the absolute signal measures in ICP-MS, which is why it is becoming a rising complementary approach to more widely established external referenced quantitative standards-based MS approaches [3]. However, it should be noted that ICP-MS is not completely reference-free, as it relies on the apriori stoichiometry of metallic heteroatom references to peptides and proteins.

In the context of quantitative immunoassays, achieving complete reference-free absolute quantification remains unattained, even with notable advancements in digital immunoassays, such as dSimoa and others, which are claimed to offer potential for effective proteoform molecule counting [4, 5]. These approaches still lack a definitive absolute signal [4], exhibit variability in performance, and have yet to achieve zero background. Designing a method of reference-free absolute quantification necessitates measures that can effectively eliminate assay backgrounds, avoiding false signals, and provide robust and precise molecular counts, referred to as the absolute signal, despite that these requirements are demanding for extremely high signal-to-noise performance. Generally, it is also essential to achieve zero background, especially in the presence of sample-dependent matrix effects, to provide also high specificity by reducing non-specific bindings. Under these circumstances, a mathematical framework can be established to accurately describe the chemical reactions and deduce absolute quantities.

### B.1 Mathematical modeling of absolute quantitative bi-component immunoassay

A minimalist model of ternary complex formation (*ternary-equilibrium-based absolute quantification model* or AQ model) was constructed in order to understand the conditions and limitations of absolute quantitative measurement regimes in bi-component immunoassays. The model describes the equilibrium formation of the assay signal-bearing ternary complex, composed of two monovalent binding antibodies bound to a target. Similar ternary-complex models were described in a range of studies investigating receptor signaling, enzymatic, scaffolding interactions and multivalent ligand-receptor binding in previous studies [10, 34–38], however, our analysis is specific for solution phase ternary immunoassays and reveals implications for pioneering absolute quantitative measurements.

Assuming that the antibody and antigen proteins are fully dispersed in solution, there is no positive or negative cooperativity between molecular interactions, and the antibodies are binding monovalently, while epitopes are sterically accessible, there is no background reaction and all couplexes are formed, the following model was devised. Let *ξ* represent the couplex concentration of the ternary complex formed by an antigen molecule with two bound antibodies. The initial concentrations of the two different antibodies are denoted by *b*_01_ and *b*_02_, while the initial antigen concentration is represented by *a*_0_. Additionally, let’s denote concentration of antigen molecules with only one antibody bound as *c*_1_ and *c*_2_, free antigen molecules as *a*, and unbound antibody molecules as *b*_1_ and *b*_2_.

By applying mass conservation

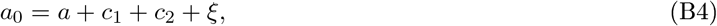

also

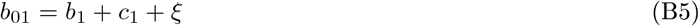

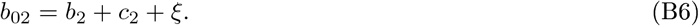

At equilibrium, mass action equations are devised, where the ratio of the concentration of the bound and unbound molecules are given by the respective dissociation constant of antibody, as

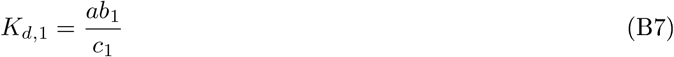

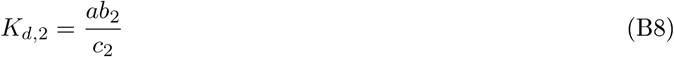

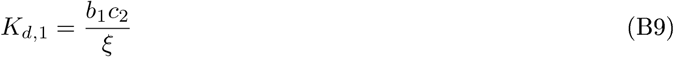

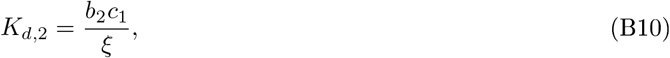

where the *K*_*d*_-s are the dissociation constants of the respective antibodies, denoted as *K*_*d*,1_ and *K*_*d*,2_, respectively.

Solving the system of equations for *a*_0_ and substituting the measured values of *b*_01_, *b*_02_ and *ξ*, and also the antibody parameters *K*_*d*,1_ and *K*_*d*,2_ gives the absolute concentration of antigen. For convenience, let us introduce the shorthand *p* = (*b*_01_, *b*_02_, *ξ*) for the measured parameters. The system of equations (B4)–(B10) has two *a*_0_ roots (non-bijective model) denoted by them as 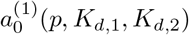 and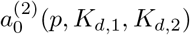, with 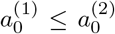 by convention, see Fig B2 for numerical modeling.

The solutions 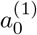 and 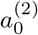 are calculated explicitly as follows:

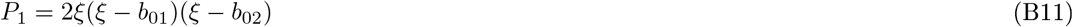

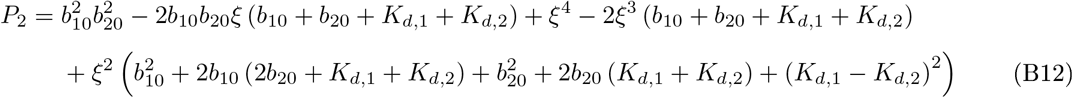

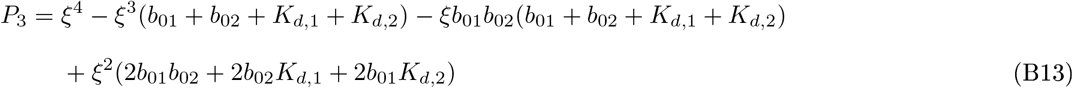

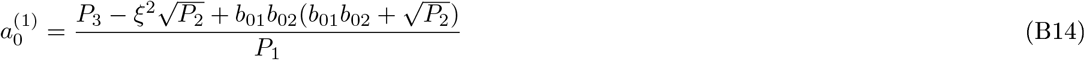

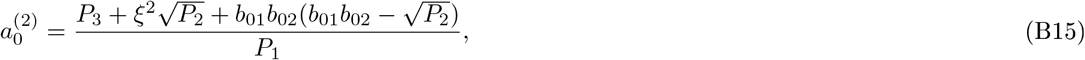

where *P*_1_, *P*_2_, *P*_3_ are auxiliary variables without any physical meaning, and are only used to simplify writing the formulae.

Yang and Hlavacek [10] formulated a mathematical framework for modeling the scaffold-mediated nucleation of protein signaling complexes, with an emphasis on quantitative determination of the formation of ternary (and higher-order) complexes. In contrast, our present mathematical formulation offers a way to calculate *a*_0_ from the measured parameters *p* in a quantitative external reference-free manner. Yang and Hlavacek [10] also showed that the equilibrium concentration of the couplex molecules *C*_2_ = *ξ* can be written as

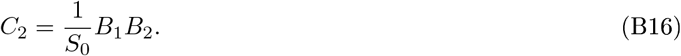

They solve (their version of) equations (B5)–(B10) for *B*_*i*_

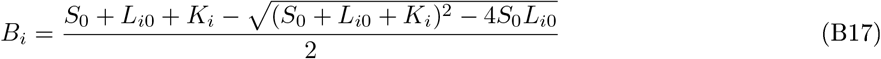

(since *B*_*i*_ ≤ min(*L*_*i*0_, *S*_0_) are physical constraints). Cited here for completeness.

### B.2 Analysis of the absolute quantitative model

The AQ model yields a non-bijective curve, as shown in Fig B2 (log-log representation), which depicts the relationship between antigen and couplex concentration. This curve comprises two distinct regions: the low antigen side (left) and the high antigen side (right), see also in Fig.1.c. On the low side, where antigen concentrations are lower than antibody concentrations, increasing antigen concentration leads to a proportional increase in couplex formation. This region exhibits a linear relationship in log-log space, culminating in the equimolar point. Conversely, on the high side, where antigen concentrations exceed antibody concentrations, rising antigen concentration competes for both antibody binding, resulting in decreasing couplex formation. The two solutions of the system of equations 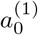 and 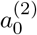 correspond to the low and high sides of the curve. The two roots define a dynamic range, typically spanning four orders of magnitude in concentrations. It’s worth noting that the region beyond the peak, the high side, is traditionally referred to as the ‘hook effect’ in the literature [39], which is considered to yield inaccurate and misleading results. However, as we have demonstrated, the hook effect is mathematically tractable.

Numerical heatmaps, focusing on the attainable LOD of the PICO assay, were also constructed (Fig A1), which depict the relationships among all the experimentally critical variables, antibody, couplex, antigen concentrations, and antibody dissociation constant (*K*_*d*_). The maps are for low side only, and the solid lines represent the ratio values between the antibody and couplexes concentrations limited by the dPCR hardwares at different *λ*s, more specifically, by the available compartment sizes (n), *λ* and analytical (couplex) LODs (*σ*, see details in caption). These lines represent the LOD conditions with attainable measurements lying to the right of the lines. Regarding the concentration of antibodies, increasing the antibody concentration results in a higher and more abrupt equimolar peak and upward curve shift, as shown in Fig. B2. Correspondingly, in Fig. B3, the iso-concentration (isomolar) lines of the antigen indicate higher concentration of couplexes until the saturation, from where couplex concentration is constant and equals the antigen concentration (i.e. saturated), also see Fig B2. The heatmaps (Fig A1) show that the highest sensitivity is slightly below of this saturating antibody concentration (i.e. the LOD line crosses the lowest ismolar). Consequently, the antibody concentration affects the attainable sensitivity, as demonstrated by the varied antigen concentrations along the LOD lines.

**Figure B2:**
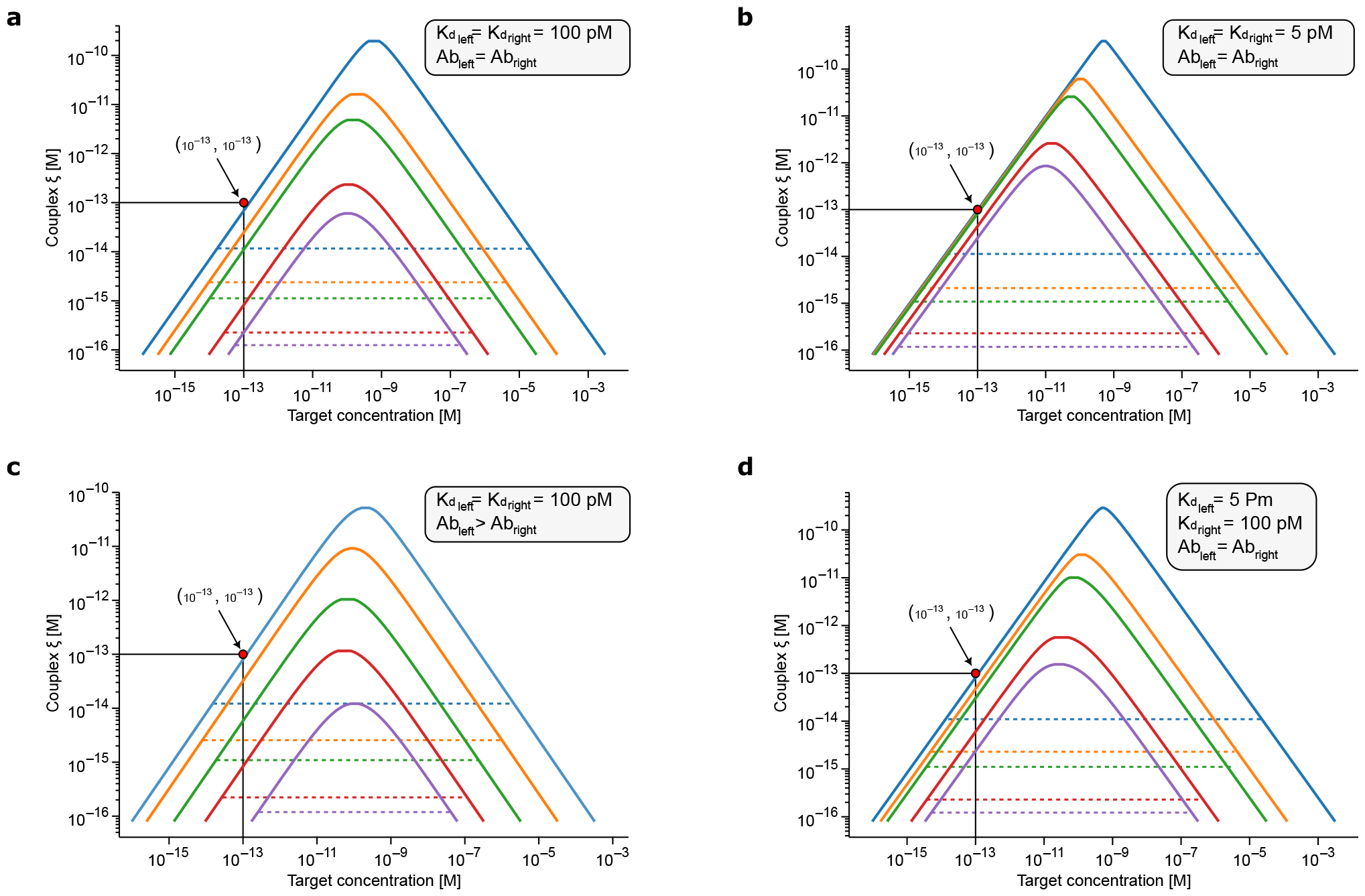
Numerical modeling the non-bijective of *p* = (*b*_01_, *b*_02_, *ξ*), depicting scenarios of different concentrations and *K*_*d*_ values of antibodies (5 and 100 pM), log-log representation. The concentration of antibodies (*b*_01_, *b*_02_) are from top to down for the curves of subfigures of (a), (b), and (d) were 500, 100, 50 10 and 5 pM (blue, orange, green, red, purple), for both antibodies, respectively (i.e. equimolar conditions), except for (c), where one of the antibodies has a concentration of 5 times less (i.e. unequimolar). The dotted lines indicate the attainable LOD assuming 0.2 million of partitions (roughly the partition size of SIMOA [11], or 8 wells of dPCR instrument Qiacuity QIAGEN (Germany, Hilden) or two 100,000 partitions, High Resolution nanowell plates of Digital LightCycler^®^ System (Roche, Switzerland, Basel)), at *λ* = 0.6 and the dPCR limit of detection of the couplex as 3 couplexes see Fig 1.b. By definition, at saturation, the curves show a 1-to-1 numerical correspondence on both axes, and the highest sensitivity curve is with the leftmost starting LOD line, also curves have a unity of slope outside of the peak region (see text, and also relative quantification B.2). **a**. Under equimolar conditions and the *K*_*d*_s are 100 pM, saturation can be achieved at 500 pM (blue) of *b*_01_, *b*_02_, while the highest sensitivity of 10.6 fM is at 100 pM (orange). **b**. Under equimolar conditions with a *K*_*d*_ of 5 pM, the curves display evident saturation for the blue, orange, and green curves, each having corresponding LODs of 13.1 fM, 2.9 fM, and 1.5 fM, respectively. Nevertheless, the curve labeled purple exhibits the highest sensitivity at 0.52 fM. **c**. Under unequimolar condition, and antibodies *K*_*d*_s of 100 pM, the highest sensitivity of 15.9 fM is the orange curve, however no saturation was reached. **d**. Equimolar condition however with two different *K*_*d*_s of 5 pM, and 100 pM, respectively, blue curve shows saturation effect with a LOD value of 15.6 fM, while the highest sensitivity is the red curve having a value of 4.2 fM. Numerically simulating conditions similar to SIMOA shows comparable sensitivities for a wide range of PICO conditions, however PICO assays are diluted and use 15 times less material. Note that as PICO has absolute signal, assuming a large enough partition size, the assay limited only by the lowest measurable number of couplexes (LOD of dPCR is 3). According to that, the maximum theoretically achievable sensitivity using an assay volume of 1 *µ*L is a persnickety 5 aM.

When comparing heatmaps with different *K*_*d*_ values, they exhibit identical saturated iso-couplex lines, indicating that the antibody’s dissociation constant (*K*_*d*_) plays a diminished role in defining antigen saturation, and only the saturation concentration of antibodies varies as a function of *K*_*d*_. This implies the potential for an absolute quantitative immunoassay, where the need to determine the *K*_*d*_ (dissociation constant) values of antibodies could be circumvented. However, achieving this would necessitate an ideal scenario: a lossless, zero background immunoassay (see main text).

To further substantiate these assertions, we investigated the mathematical characteristics inherent in the 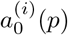 model. Based on Figs B2 and B3, our hypothesis posited that variations in *K*_*d*_ values primarily impact the low side of the curve, while different antibody concentrations affect both sides. Furthermore, it is evident that at saturation, the low side of the curve is independent of the specific *K*_*d*_ values. To investigate this proposition, we examined the derivative of *a*_0_ with respect to *K*_*d*_ and antibody concentrations. To simplify the visualization, we assume for the rest of this subsection that *K*_*d*,1_ = *K*_*d*,2_ and *b*_01_ = *b*_02_.

To motivate our analysis, recall that the rate of change can be approximated by the derivative formula:

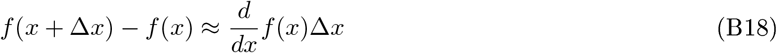

for a continuously differentiable function *f* with *x* ∈ ℝ and Δ*x >* 0. Equation (B18) suggests that we can analyze the rate of change, that is, the sensitivity of 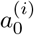 to the different parameters by looking at the logarithm of the model (with the model 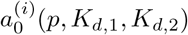 as *f*). Fig B4.a shows the derivative of log *a*_0_ with respect to log *K*_*d*_. The figure indicates that sensitivity to *K*_*d*_ is asymptotically constant as the couplex concentration is decreased in the experiments. This means that the (log) sensitivity rate is constant across the side of the *ξ*-*a*_0_ curve, with deviations in the peak regions (curved ends of the horizontal lines in Fig B4.a). Constant sensitivity rate appears as parallel (sides of the) curves in Fig A1. The figure also shows that this constant derivative goes to 0 as the *K*_*d*_ is decreased relative to the antibody concentration. Note that the (log) derivative being 0 implies the measurement is not sensitive to the *K*_*d*_ value used in the calculations of antigen concentration, that is, the dissociation value does not affect the outcome of the experiments.

In a similar manner, Fig B4.b shows the derivative of log *a*_0_ with respect to log *K*_*d*_. The same asymptotically constant behavior can be observed at low couplex concentrations, that is, at the side of the *ξ*-*a*_0_ curve.

These findings imply the viability of an absolute quantitative assay relying exclusively on the 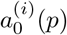 model, while strategically substituting a surrogate *K*_*d*_ value into the equations, taking into account an optimal precision around the peak region.

To grasp the precision implications and the influence of introducing an arbitrary surrogate *K*_*d*_ value into the equations, selected to enhance precision, we simulated a logarithmic difference between the default values and those computed using the *p* = (*b*_01_, *b*_02_, *ξ*) model. To quantify, we conducted simulations with various surrogate *K*_*d*_ values to assess their impact on the precision of absolute quantification. The colorbar shows the log_10_ difference between the actual value and the model-predicted absolute amount, that is, a value of 0.3 represents a twofold difference. Observations indicate that, as expected due to the absence of *K*_*d*_ dependence, the precision on the high side remains consistently high. On the low side, precision remains high until the actual value *K*_*d*_ value is below 100 pM, with this trend holding for all surrogate *K*_*d*_ values above 10 pM. This assessment suggests that an antibody concentration of 500 pM is sufficiently high to provide accurate absolute quantities with true *K*_*d*_ values lower than 100 pM. However, importantly, the saturation concentrations can be experimentally determined, even the (effective) *K*_*d*_s of antibodies can be estimated. The experimental strategy employed is maintaining a constant antigen concentration while systematically varying the antibody concentration, referred to as isomolar titration (IT) Fig C7.c (see details there).

Minimizing the variance of *a*_0_ between isomolar experiments, while varying the *K*_*d*_, can serve as a useful method for estimating this effective dissociation constants (*K*_*d*_) of the antibodies, see C7. This approach yields an effective *K*_*d*_ value, as it takes into account other experimental variables (see B.4). While this approach is theoretically viable, we have observed its sensitivity to experimental conditions. Nonetheless, it presents a promising avenue for further experimental evaluation to enhance the applicability and reliability of absolute quantitative PICO analyses.

### B.3 Relative quantification

Besides absolute quantification, relative quantification is also possible and leverages the observed linear log-log behavior with a slope of unity (m =1). As per definition,

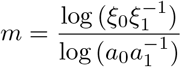

where *ξ* is the concentration of couplexes of relative measurements, and *a* is the corresponding antigen concentrations, which leads to

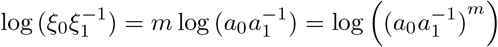

Notice that 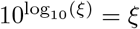, therefore, the logs can be inverted to

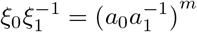

where, as it was shown, m = 1, and it leads to

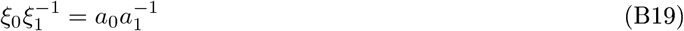

hence the ratio of the couplex concentrations or just the number couplex counts equals the ratio of absolute analyte concentrations. Nevertheless, albeit couplex counts are also applicable, to mitigate potential experimental biases, it is advisable to apply corrections such as *λ*-normalization and no-analyte control (ABC - antibody control) correction. However, the application of labeling Δ correction is optional, as it merely represents a linear transformation that does not impact the ratio.

With relative quantification the concentrations of antibodies employed become irrelevant, as the ratios of the couplexes concentrations at all antibody concentrations yield molar ratios suitable for relative comparisons, thereby eliminating the necessity for isomolar titration. It is acknowledged that the peak region of the curve is unsuitable for relative quantification analysis due to its significant deviation from linear behavior. Nevertheless, the versatility of employing relative quantification across a broad spectrum of antibody concentrations renders it favorable in selected applications.

### B.4 Effects of experimental conditions on the absolute quantitative model

In contrast to the idealized model, which assumes perfect antibody quality and labeling, we have taken into account various sources of errors to assess their impact on measurement precision. These sources include the proportion of labeled antibodies (*β*), label synthesis errors that render the label undetectable (*γ*), the active fraction of antibodies capable of binding (*α*), and the presence of free labels (*ϵ*).

Let *α*_*i*_ represent the fraction of active antibody molecules for antibody *i* (e.g., if the concentration of the antibody molecules is *b*_01_, then the concentration of active antibodies is *αb*_01_).

We can incorporate *α* into the model by modifying equations (B7)–(B10) as follows:

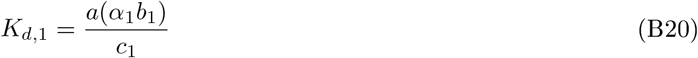

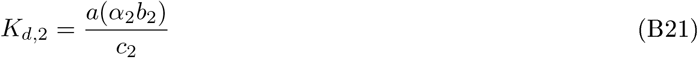

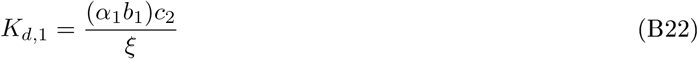

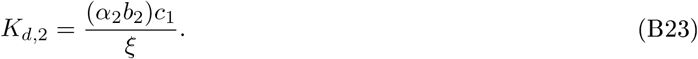

By rearranging equations (B20)–(B23), we obtain the following forms:

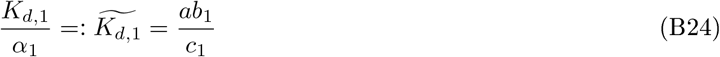

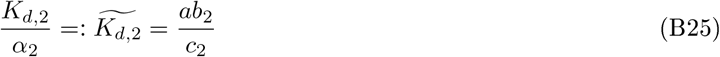

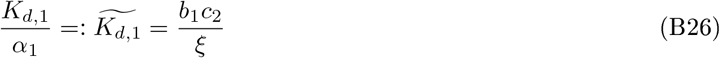

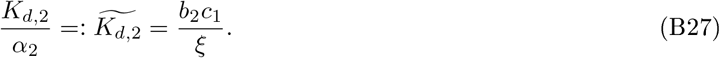

*α*(≠ 1) can be conflated with the dissociation constants, resulting in a new, effective, *K*_*d*_ value. Remarkably, in the saturation regime, where the analyte concentration *a*_0_ becomes independent of the antibodies’ dissociation constants (*K*_*d*_), the need to measure *α* is likewise eliminated. Considering the inherent challenges associated with determining *α*, this fortuitous simplification significantly enhances the reliability of the assay.

It can be demonstrated that the presence of free labels (*ϵ*) can be handled similarly. The elimination of determination *ϵ* is also desired, as it is challenging to measure, and reinforcing the model’s robustness and reliability under practical conditions, where some extent of free labels in the antibody preparations is unavoidable. Note that a high level of free labels necessitates a higher dilution to achieve the same dPCR *λ*, which inevitably leads to lower sensitivity of the assay.

The proportion of labeled antibodies is an independent parameter within the model *p* = (*b*_01_, *b*_02_, *ξ*), and it can be incorporated in the model as follows:

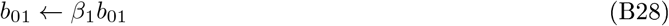

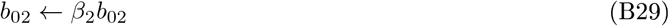

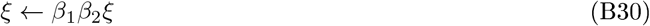

where *β*_1_ and *β*_2_ denote the proportion of labeled antibodies for the two antibodies, respectively. It is important to note that if the precise values of *β*_1_ and *β*_2_ are known, it becomes straightforward to reverse this transformation and recover the chemically active antibody concentrations. It also can be shown that no transformation is possible to conflate *β* parameters to the effective *K*_*d*_, consequently the proportion of the labeled antibodies need to be independently determined. To get the chemically active concentration of the antibodies and the couplexes the reverse transformation, labeling normalization, is applied, D.5). In this study, the *β*s and corresponding Δ were determined by protein SDS-PAGE (see D.3).

The label error (*γ*), resulting from synthesis failures that generate undetectable label sequences, has identical analytical effects as the proportion of labeled antibodies (*β*) and can be mathematically described in the same form. In other words, antibodies with label errors can be considered as antibodies that are unlabeled. These errors are typically minimal when using contemporary oligonucleotide chemical synthesis methods and are often deemed negligible. However, *γ* needs to be taken into account and corrected for in highly sensitive measurements.

We have also investigated the impact of cooperativity between antibodies, a topic previously discussed in relation to a similar model [10]. Typically, low cooperativity in antibody bindings is expected, consistent with the published lack of cooperativity between rituximab and ofatumumab [40](Weiner, 2010). For instance, rituximab’s epitope spans amino acid residues 168-175 of the CD20 protein’s large extracellular loop, while ofatumumab epitope involves both the large and small CD20 extracellular loops, the two epitopes are separated by only six amino acids, the non-cooperative binding of the antibodies was also confirmed using PICO (unpublished). Similarly, our poli-tag rTRXpeptide data demonstrate the absence of cooperativity C6 and 1. Nevertheless cooperativity is expected in immunoassays, particularly negative cooperativity or blocking, and this aspect has been overlooked in other bi-competent immunoassays, such as capture ELISA. PICO possesses the unique capability to detect cooperativity through the triangular measurement concept, as seen above. Significantly, in an unpublished study, we investigated two antibodies targeting the same 4EBP1 epitope — one specific to its phosphorylated state and the other to its unphosphorylated state. This study confirmed a perfect mutually exclusive cooperativity, selectively determined by the phosphorylation status of the epitope, as anticipated.

**Figure B3:**
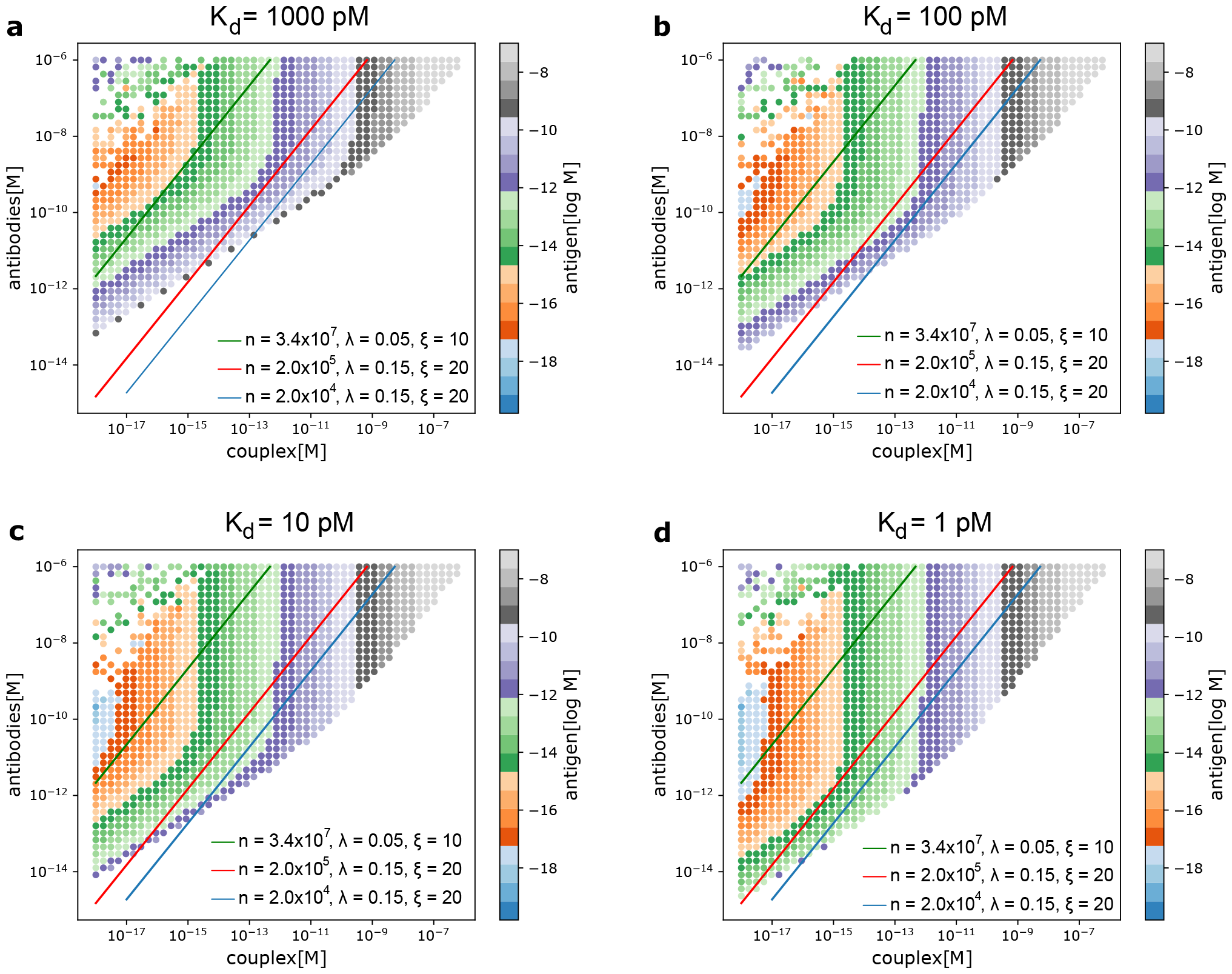
Simulation of the attainable sensitivity of PICO. The logarithm of *K*_*d*_s are -9 (a), -10 (b), -11 (c), and -12 (d), respectively. Some combinations of values calculate physically impossible complex roots or stoichiometrically impossible scenarios, where e.g. the concentration of couplexes exceeds that of antibodies, these indicated as white areas. Solid lines represent LODs determined by dPCR hardwares, with attainable measurements to the right of these lines, and represent the attainable signal-to-noise (S/N) ratios in dPCR. Parameters include the compartment size (n) at *λ* (the average number of antibodies per compartment, defining the number of antibodies) and analytical (couplex) LODs (*σ*) (simulated data, see Fig.A1). The blue line corresponds to n = 25,000 at *λ* = 0.15 and *σ* = 20 couplexes, resulting in a ratio of 187.5 (Qiacuity, Stilla). The red line corresponds to n = 200,000 at *λ* = 0.15 and *σ* = 20 couplexes, yielding a ratio of 1,500 (roughly corresponds to SIMOA conditions). The green line corresponds to n = 34 millions at *λ* = 0.05 and *σ* = 10 couplexes, resulting in a ratio of 215,000 (largest published dPCR compartment size [12], Enumerix). All ratios are calculated using the formula, S/N = *c* × *λ/σ*. All in molar units. Above the saturation concentration (see the vertical part of the isomolar lines of antigen), the couplex concentration remains constant and equals the antigen concentration, regardless of *K*_*d*_ (all figures are congruent), demonstrating *K*_*d*_ independence in saturation. However, the saturation concentration is *K*_*d*_-dependent, with lower *K*_*d*_ values resulting in lower saturation concentrations (see the start of the vertical isomolar). Lower compartment sizes cover the low picomolar to medium femtomolar regimes, while higher-end conditions encompass sensitivities from low femtomolar to low attomolar ranges, setting the theoretical maximum (i.e. 5 aM).

**Figure B4:**
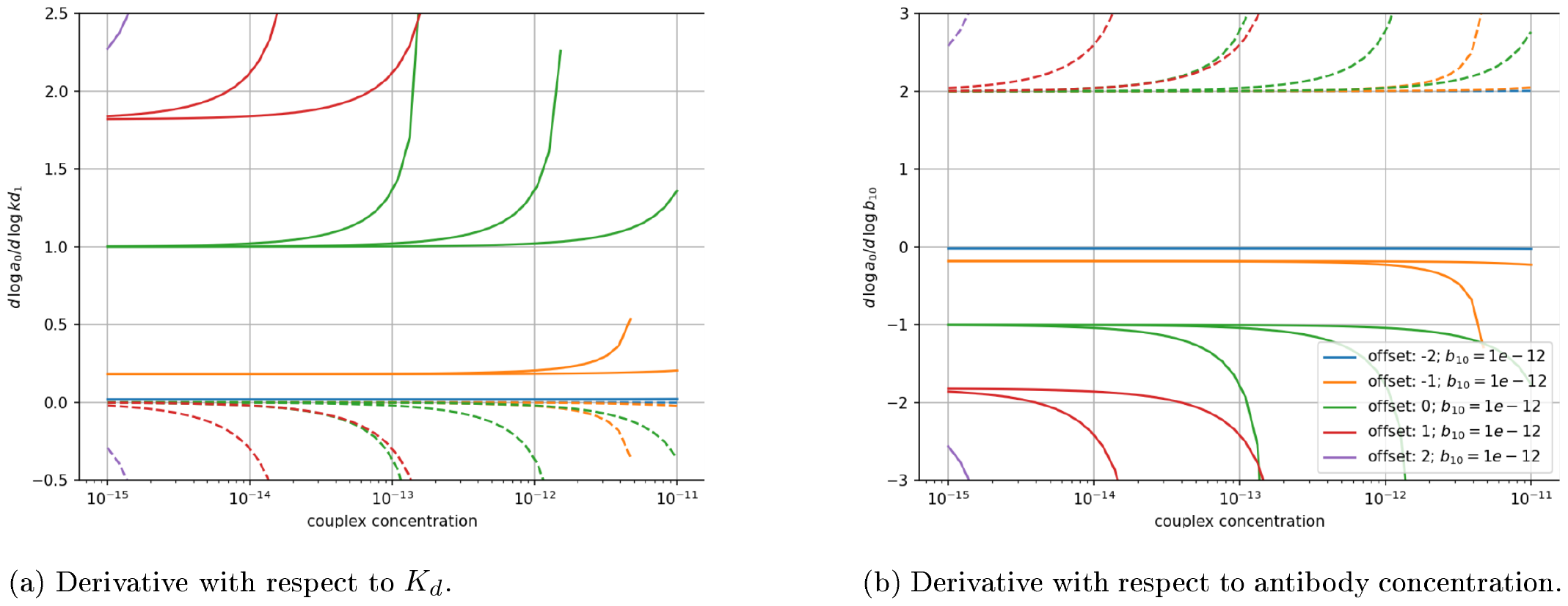
Logarithmic derivatives derivatives of log *a*_0_ with respect to the logarithm of the antibody dissociation constant, *K*_*d*_ (a) and concentration (b). Solid lines represent the low side of the curve, while dashed lines depict the high side. Logarithmic offsets are indicated by colors, the log_10_ difference between *K*_*d*_ and antibody concentrations (that is, log_10_ *K*_*d*,1_ − log_10_ *b*_01_), where the latter is set at 1 pM, and *K*_*d*,1_ = *K*_*d*,2_ and *b*_01_ = *b*_02_ assumed. As shown in **a**., the low side of the curve shows *K*_*d*_ dependence of *a*_0_ (derivative is higher than 0), but this effect diminishes as the antibody molar concentration surpasses the *K*_*d*_ value, becoming low at offset = -1 (10 times higher antibody concentration than *K*_*d*_), or negligible at offset = -2 (100 times higher). Conversely, the high side of the curve remains entirely unaffected by the *K*_*d*_ values. Both sides exhibit deviations from this behavior in the peak region of the curve. Regarding **b**., the low side of the curve at offset = -2 exhibits characteristic saturation behavior, demonstrating independence of *a*_0_ from antibody concentration. In contrast, the upper portion of the curve displays consistent (nonzero) dependence regardless of the offset. This insight motivates the development of an assay based on the 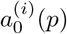 model at saturation. Such an assay would eliminate the need to determine *K*_*d*_ values, as it enables the *K*_*d*_-independent determination of *a*_0_ and simplifies the assay by relying on easily measured antibody concentrations instead.

**Figure B5:**
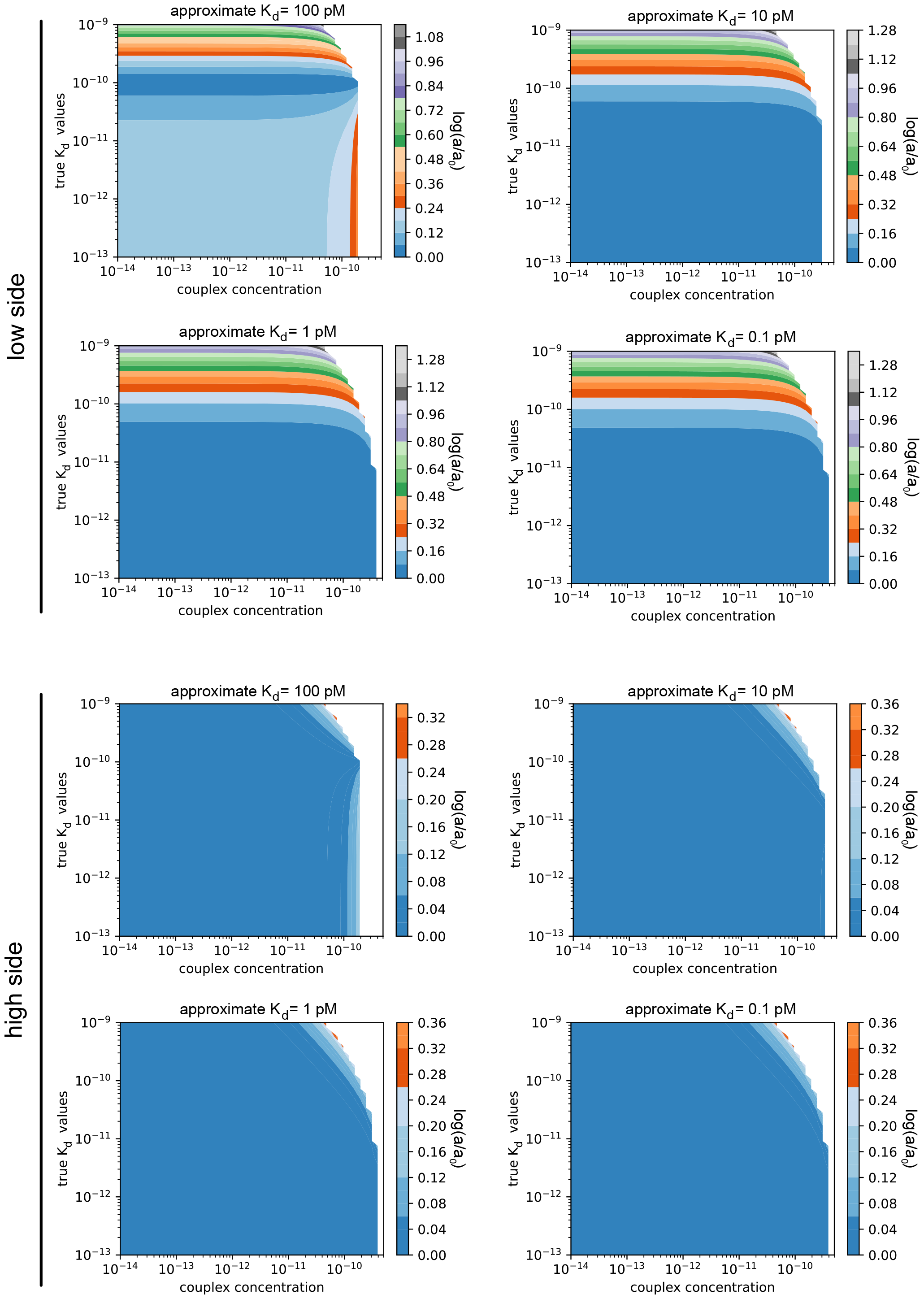
Relative precision of absolute concentration of analyte assuming 5.0 × 10^−10^ M antibody concentration while the surrogate *K*_*d*_ of antibodies are varied. X-axis is the concentration of couplexes, while Y-axis represents the true *K*_*d*_ value. The color coded values are the logarithm of the ratio of the calculated and real *a*_0_. The upper graphs show the low side while the bottom graph represent the high side of the curve, log_10_(2) = 0.3, which indicates the two-fold difference, was taken as an precision threshold. Based on this analysis, it is evident that the choice of a surrogate *K*_*d*_ is flexible, provided it exceeds the saturation concentration of the given antibody. When using a surrogate *K*_*d*_ of 10 pM or lower (also a safe default is 50 pM), the low-side exhibits remarkable precision, particularly when the true *K*_*d*_ falls below 200 pM. It’s noteworthy that the peak region, situated at the right edge of the graph and neighbored by a mathematically inapplicable white area, maintains precision akin to the entire curve on the low side or only nuancely influenced by the couplex concentration on the high side. Therefore, using the the model of *p* = (*b*_01_, *b*_02_, *ξ*) even the peak region attains high precision using a proper surrogate *K*_*d*_, however to avoid *K*_*d*_ above the threshold it is recommended conducting IT experiments to ensure high precision.

## Appendix C Additional figures

Further validations.

**Figure C6:**
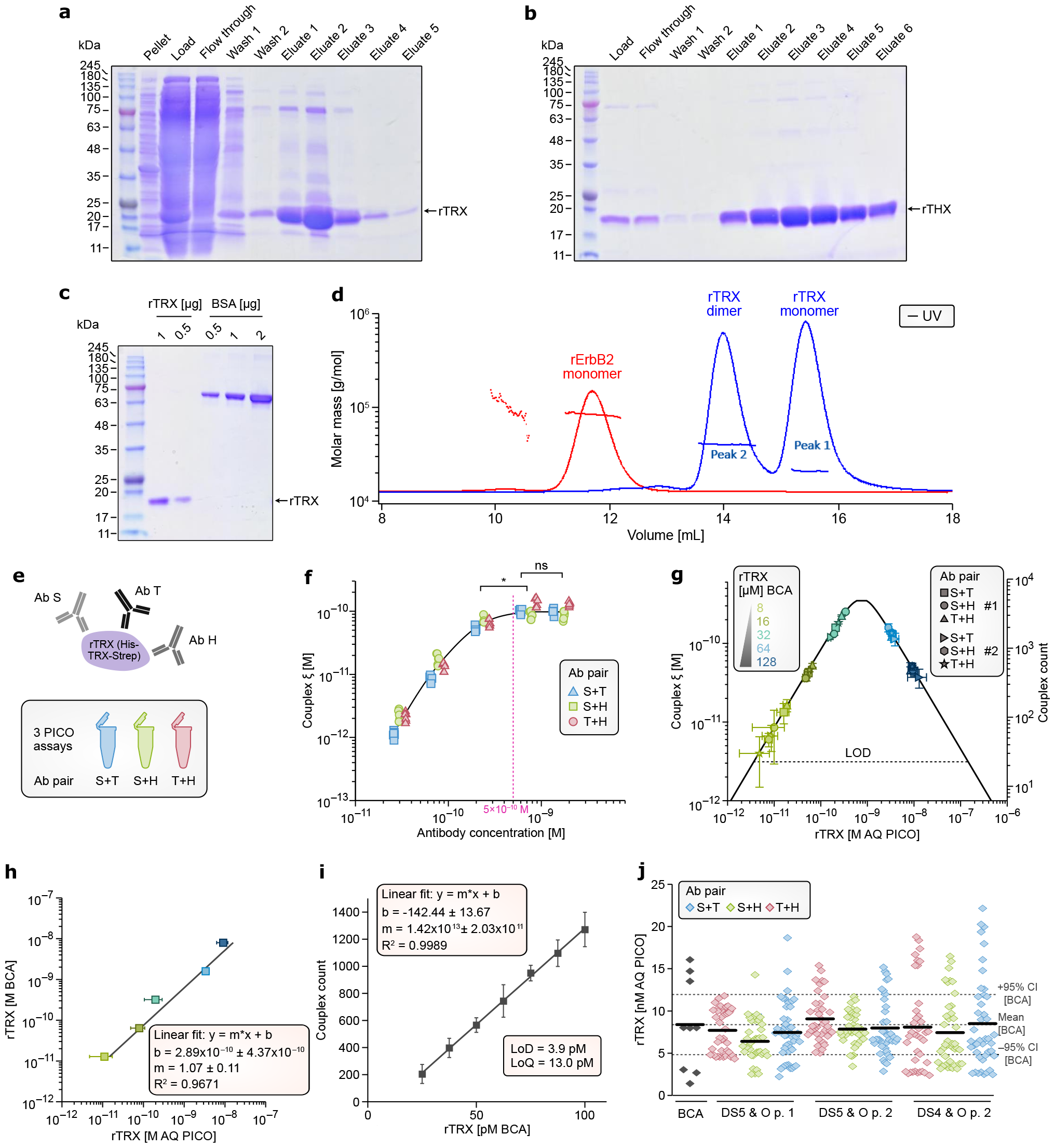
Purification of rTRX D.1. SDS-PAGE of Ni-NTA (a) and STREP-Tactin (b) fractions. **a**. lanes are left to rigth E. coli pellet, the lysed sample (load), flow-through (FT), wash (W1 – W2) and fractioned eluates (E1-E5), while **b**. pooled Ni-NTA elutes as load, flow-through (FT), wash (W1 – W2) and fractioned eluates (E1-E6) **c**. The purity of rTRX compared to BSA. **d**. SEC-MALS results of the rTRX and rErbB2, depicted elution volume vs molar mass (overlay of rTRX and rErbB2). The molecular weights of rTRX was 20.8 kDa and for rErbB2 82.6 kDa were determined, rErbB2 contains 3.8% (m/m) of higher MW impurity, unlikely being ErbB2 aggregates, ErbB2 is considered to be pure with molarity corrected for the impurity, while rTRX partially forms dimers, where the 52% of mass is in dimer form, and it is assumed that it reduces the m/v concentration based dPCR countable molarity of rTRX by 34.2%, which compensated in the final AQ PICO results. Absolute quantification of recombinant thioredoxin. **e**. Schematic representation of the triangular PICO assay. The BCA quantified rTRX was reacted with anti-penta-his (H), anti-thioredoxin (T), and anti-strep-tag-II (S) antibodies, respectively see D.1. **f**. Isomolar titration, determination of the saturation concentration of the antibodies. Concetration of antibodies were from left to right of 25 pM, 75 pM, 220 pM, 660 pM and 2 nM, respectively - and PICO binding reactions were set up using of a constant, (isomolar) concentration of rTRX of 200 pM (see D.5). The molar concentration of the formed couplexes (*ξ*) was determined and showed saturation by the increasing ABX concentration. The minimal saturating concentration was determined by ANOVA (Kruskal-Wallis ANOVA with Tukey’s test, *p < 0.05, n=90) being less than 500pM. Different markers indicate the different ABPs (see legend). **g**. AQ (calibration) curve of recombinant rTRX. Symbols indicate different ABPs, the assay carried out using two different batches of antibodies (#1 and # 2) at 500 pM of ABX (antibody mix), each marker representing the mean of 8 replicates and with standard deviations are shown (mainly imperceptible), n=144. The dotted line denotes the LOD of measurement (20 couplexes, LOD 3.9 pM). The average AQ values for each dilution of the sample (n=48) accurately reproducing the reference concentrations measured with BCA (t-test indicated) in order from blue to green, 8.38 ± 5.45 nM BCA to 9.91 ± 2.77 nM PICO (p = 0.43), 1.68 ± 1.09 nM BCA to 3.35 ± 4.97 nM PICO (p = 0.0016), 335 ± 218 pM BCA to 196 ± 91.0 pM PICO (p = 0.09), 67 ± 43.5 pM BCA to 78.3 ± 29.6 pM PICO (p = 0.47), and 13.4 ± 8.72 pM BCA to 11 ± 6.23 pM PICO (0.45). **h**. Correlation of PICO AQ values against the BCA reference. The regression line has a slope of 1.07 ± 0.11, intercept 290 ± 440 pM and *R*^2^ = 0.99 with n = 234, confirms 1-1 lossless counting. **i**. Correlation AQ plot of couplexes and the BCA reference measuring rTRX, within close vicinity of the limit of detection (LOD). LOD was 3.94 pM and LOQ 13.02 pM, slope m = 1.42 × 10^13^ ± 2.03 × 10^11^, y-axis intersection b = -142.44 ± 13.67, R2 = 0.99, n=42. rTRX concentrations are 100 pM, 87.5 pM, 75 pM, 62.5 pM, 50.0 pM, 37.5 pM, 25.0 pM, 12.5 pM and 1 pM. **j**. The AQ of rTRX measurements combining all dilutions per ABPs each replicated under inter-assay conditions (two operators), and two different dilution series (DS4 and DS5, 4x and 5x dilution series) referenced against BCA measurements (8.38x10-9 ± 5.45x10-9 M BCA, n = 9 to 7.82x10-9 ± 3.70x10-9 M PICO, n = 363, t.test (t(8)=-0.307, p = 0.766)), the 95% confidence intervals of the BCA measurement indicated as dotted lines. ANOVA shows no significant differences between all groups (F(7, 278) = 0.669 p = 0.698, n=285).

**Figure C7:**
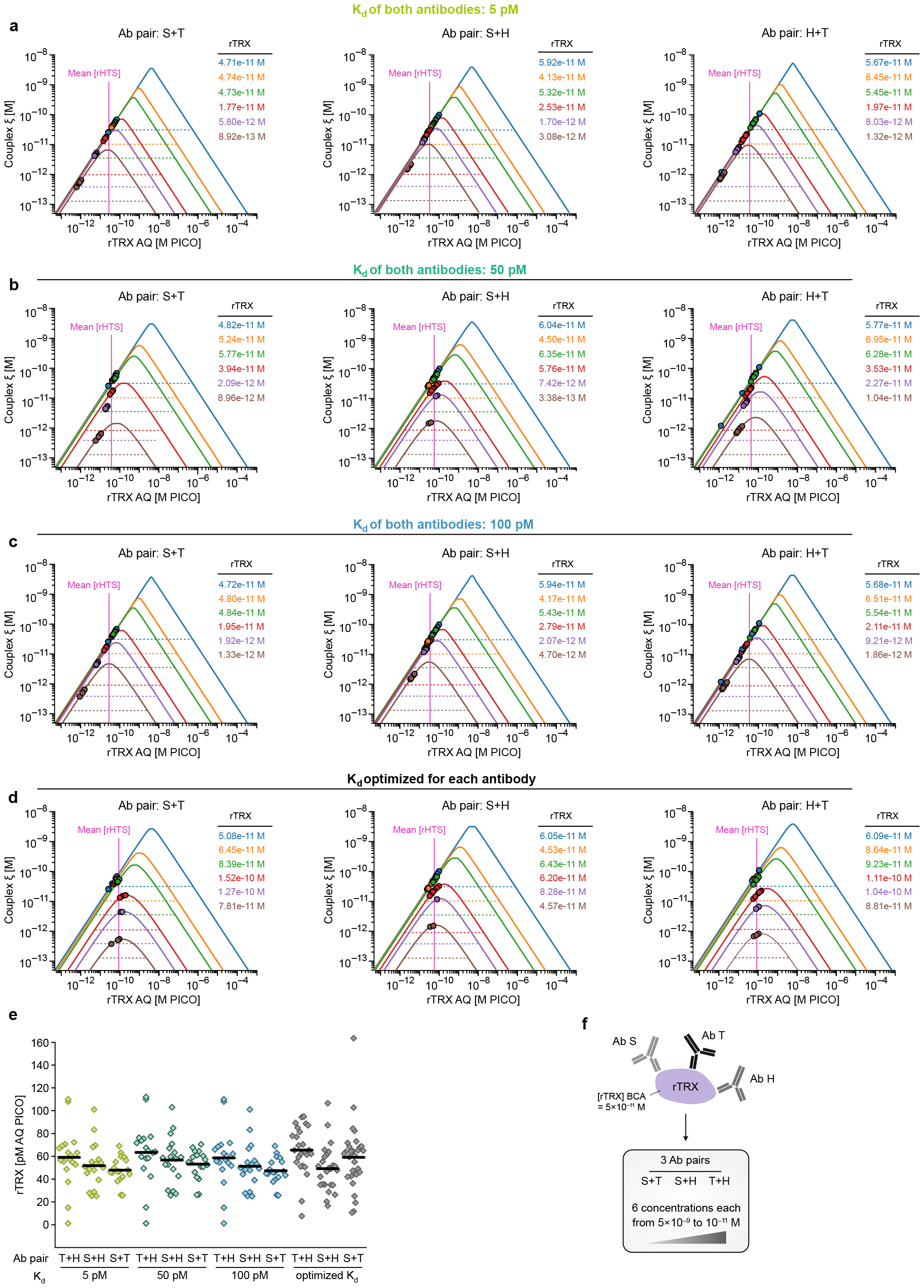
Triangular isomolar titration of rTRX. AQ curves *ξ* (observed) is plotted against AQ molar concentration of rTRX (according to*p* = (*b*_01_, *b*_02_, *ξ*) model). This titration was carried out at various antibody concentrations (ABXs): 5 nM (blue), 1 nM (orange), 500 pM (green), 100 pM (red), 50 pM (purple), and 10 pM (brown). The isomolar concentration of rTRX was 50 pM (5 × 10^−11^). Dotted lines represent the limit of detection (LOD), which varies with ABX concentration. The model-calculated AQ concentrations of rTRX are ABX color-coded for each ABP and applied surrogate *K*_*d*_ values as indicated. The mean of all model-based calculated rTRX concentrations are indicated with a vertical red line. **f**. Schematic representation of the triangular rTRX PICO assay (anti-penta-his (H), anti-thioredoxin (T), and anti-strep-tag-II (S) antibodies), see also D.1. **a,b,c** . The applied surrogate *K*_*d*_s are 5 pM, 50 pM, and 500 pM, respectively. Based on the *ξ* values obtained the 5 nM (blue), 1 nM (orange), 500 pM (green) ABX concentrations are saturated (*ξ* equals antigen concentration), and predicting rTRX AQ concentrations with minimal %SD (not shown). Lower ABX concentrations show significant departures (higher %SD) from the isomolar concentration of rTRX. **d**. Optimized *K*_*d*_ values for the antibodies. The *K*_*d*_ values were determined by minimizing the %SD of the means of rTRX for all ABPs, resulting in the following *K*_*d*_ values, anti-Strep antibody at 5 × 10^−11^ M, anti-HIS antibody at 6 × 10^−11^ M, and anti-TRX antibody at 3 × 10^−10^ M. All of the *K*_*d*_s are smaller than the saturated ABX concentrations, except for anti-TRX antibody, which is slightly higher, confirming findings regarding robust precision (see B5 and C6). By definition, all predicted rTRX concentrations are in close agreement with the isomolar concentration of rTRX. **e**. Plot representing the mean and distribution of AQ values of rTRX for different surrogate *K*_*d*_s and minimized-SD *K*_*d*_s (the only measurements with a saturated ABX concentration). No significant difference was observed (ANOVA, F(11, 229) = 1.62, p = 0.093, n = 240), confirming the robustness (the independence of the AQ model from *K*_*d*_ at saturation) of the calculation of AQ molar concentrations using different surrogate *K*_*d*_s.

**Figure C8:**
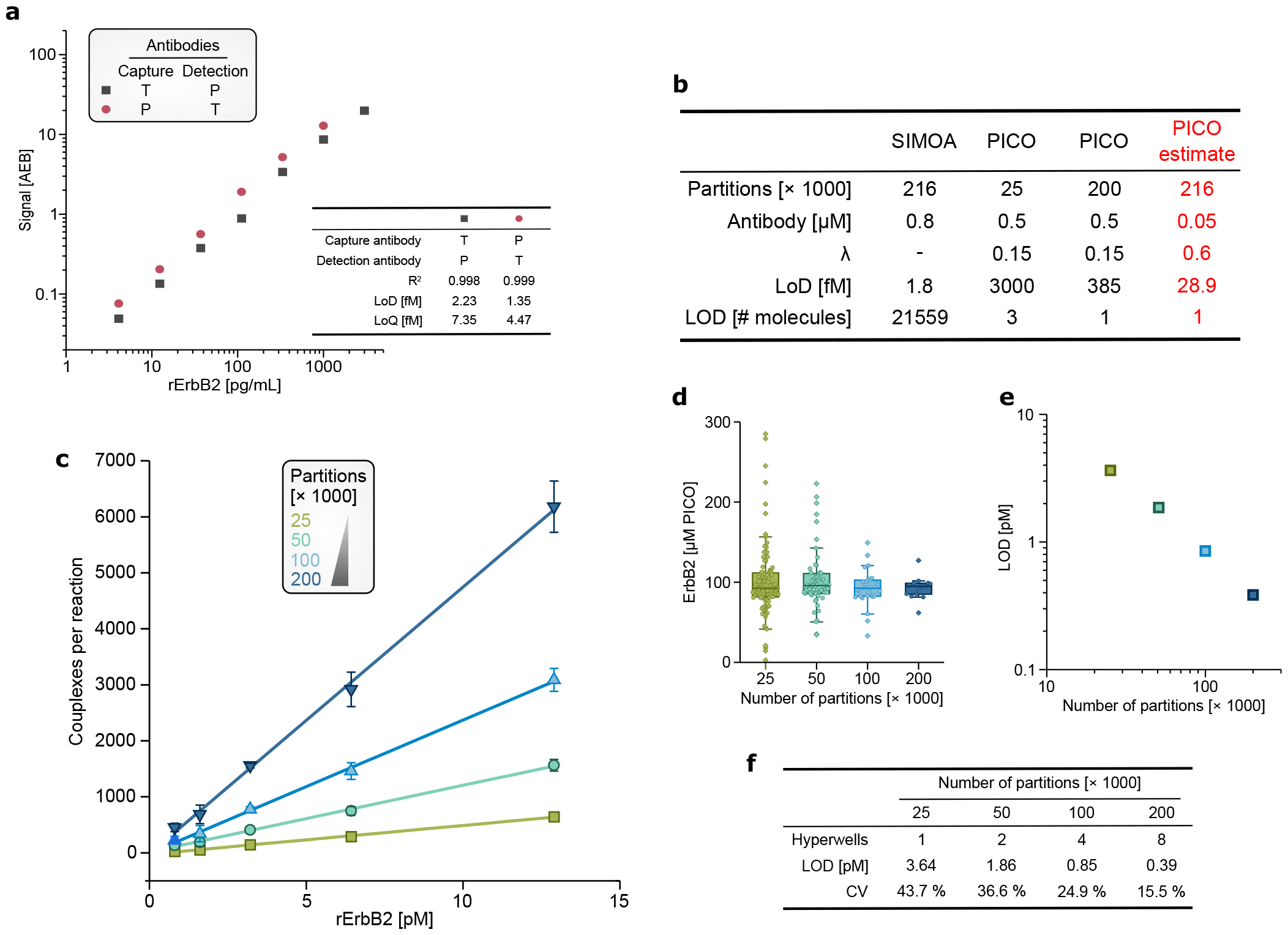
Comparing high sensitivity measurements using SIMOA and PICO. **a**. The quantification of rErbB2 using the SIMOA. Antibodies are trastuzumab (T) and pertuzumab (P) as capture and detection antibodies (varied), respectively. Measurements were 3000 pg/ml, 1000 pg/ml, 333 pg/ml, 111 pg/ml, 37 pg/ml, 12.3 pg/ml and 4.12 pg/ml. The legend table shows the linear regression and the limit of detection (LOD). **b**. The table shows the comparison of PICO and SIMOA assays. Conditions are indicated, *λ*, the measured LOD (M) and LOD (# molecules). The red font indicates extrapolated data. SIMOA demonstrates superior sensitivity at lower or comparable compartment size. However, considering sample volume, at the same compartment size, SIMOA and PICO exhibit similar sensitivities. Notably, PICO employs a high dilution before the dPCR detection step, which is reflected the vastly different LOD [molecules] (20,000 times lower for PICO). **c** .Relationship between LOD and precision of AQ concentrations of PICO as function of partitions size (hyperwelling). Concentrations of ERbB2 were 12.90 pM, 6.43 pM, 3.21 pM, 1.61 pM, and 0.80 pM, ABX is 500 pM, Qiacuity 26K nanoplates were used with 1, 2, 4, or 8 wells hyperwelled, respectively. **d**. Boxplot represents the mean and distribution of AQ values (ANOVA F(3, 202) = 0.532, p-value = 0.661, n = 205), indicating decreasing SD over increasing compartment sizes. **e**. Plot represents the decreasing concentration of LOD over increasing compartment sizes (log-log), showing a linear correlation between number of partition and LOD. **f**. The table presents values of limit of detection (LOD), LOD [molecule].

## Appendix D Materials and Methods

### D.1 Materials

PIC-PBS was made dissolving one tablet of cOmplete, EDTA-free, protease inhibitor cocktail (04693132001, Roche) in 50 ml of PBS (14040133, Gibco). ABC and LBTW buffers were from Actome (PICO AMC Kit, Actome). Pierce*™* BCA Protein Assay Kit (23225, Thermo Scientific) (protein quantification) and human ErbB2/Her2 Quantikine ELISA kit (DHER20, R&D Systems) were used according to the manufacturer’s instructions. Recombinant HER2 (rErbB2) (10126-ER, 70kD extracellular domain, R&D Systems) was dissolved in LBTW, stock concentration of 0.04 μg/μl, incubated for 15 min at 30 °C and sonicated for 5 min, aliquoted, and stored at -20 °C until use, before use it was diluted in ABC buffer. Recombinant STREP-thioredoxin-HIS (rTRX), full peptide sequence is N’-MASA-<u>WSHPQFEK</u>-iSRELVDPNSQiSGSSDKiiHLTDDSFDTDVLKADGAiLVDFWAEWCGPCKMIAPiLDEIADEYQGKLTVA-KLNiDQNPGTAPKYGiRGiPTLLLFKNGEVAATKVGALSKGQLKEFLDANLAGSACELGTPGRPAAKLA-AAQLYTRASQPELAPEDPEDLEHHH-<u>HHHHH</u>-C’ (STREP and HIS tags are underlined) was expressed in E. coli. Briefly, the codon optimized sequence without the tags was cloned into His-Strep pQE-TriSystem vector (33903, QIAGEN) and expressed in E. coli BL21 DE3 (genotype: F– ompT hsdSB (rB–, mB–) gal dcm (DE3)). Cells were lysed by repeated freeze-thaw cycles and lysis was tested by SDS-PAGE samples of pelleted E. coli and supernatant. Cell lysis buffer contained 50 mM NaH_2_PO_4_, 300 mM NaCl, 10 mM imidazole at pH 8. The supernatant of clarified extract was loaded to Ni-NTA (30210, QIAGEN) resin and gravity flow affinity chromatography was applied. Binding was allowed for 60 min. Resin was washed twice with lysis buffer containing 30 mM imidazole and eluted with lysis buffer containing 250 mM imidazole. Elution fractions were tested by SDS-PAGE analysis for eluted protein (C6.a) and pooled for further purification via Strep-Tactin (30004, Qi-AGEN). The pooled eluate was STREP-Tactin resin purified similarly, expect using STREP wash buffer (lysis buffer without imidazole) and the elution was performed at wash buffer plus 2.5 mM Desthiobiotin. Successful purification was validated via SDS-PAGE (C6.c) using Serva Triple Color Protein Standard III (39258.01). The quality of rTRX was validated using SEC-MALS and HPLC (C6.e.). Briefly, size-exclusion chromatography with multiangle light scattering (SEC-MALS) was done using HPLC (1260 infinity ii LC System, Agilent) with Citivia Superdex 200 increase 10/300 GL column applying 40 μg TRX in PBS pH 7.4 at 0.75 ml/min detected with UV280 and MALS device (miniDAWN, Waters Wyatt), which analyzes the molecular weight of protein samples according to two dimensions with a size-exclusion chromatography by volume and scattering by molecular weight (MW). Data were analyzed using Astra 8.2.0 software. rErbB2 was also analyzed with SEC-MALS.

### D.2 Processing cells for the PICO assay

MCF7 (HTB-22TM, ATCC) and BT-474 (HTB-20TM, ATCC) cells were cultured under standard conditions. The cell were untreated, or where indicated, dactolisib treated. Briefly cells were washed twice with PBS and then incubated at 5.6 μM dactolisib (ab120882, Abcam) for 4 hours in culture medium, then cell culture medium was removed, and the cells were washed twice with PBS. The mock-treated also washed twice with PBS. The cells were harvested and washed in PIC-PBS two times (400x g for 5 min), and counted (Countess, Invitrogen). After pelleting at 400x g for 5 min, one million cells were lysed in 100 μl LBTW lysis buffer. The lysis was carried out for 3 h at 4 °C, sonicated for 5 min and homogenized using a QIAshredder (20000x g for 2 min) (79656, QIAGEN) ensuring a molecularly-disperse protein sample. The lysate was further diluted in LBTW as indicated.

### D.3 Antibodies and labeling

Pertuzumab (PTZ, anti-ErbB2, Roche), trastuzumab (TTZ, anti-ErbB2, Roche), anti-ErbB3 2F12 (MA5-12675, Invitrogen), anti-ErbB3 Ab90 (MA5-12867, Invitrogen), penta-HIS (34660, QIAGEN), anti-TRX (4C12H10, antikoerper-online), recombinant anti-STREP II (11A7, Abcam), GAPDH (1E6D9, Proteintech), anti-4EBP1 554 (AHO1382, Invitrogen), anti-4EBP1 4F3 (H00001978-M01; Abnova), anti-4EBP1 phosphorylation T46 (MA5-27999, Invitrogen), anti-S6K1 2C2 (SAB1412617, Sigma-Aldrich), and anti-S6K1 (609902, Biolegends) antibodies were used as indicated. PICO detector antibodies were labeled in a site-, and heavy chain specific way with amplifiable DNA oligonucleotide labels (PICOglue Antibody Labeling Kit and PICOglue BL, P8, N6, O7 Label Kits, Actome). In brief, the antibodies were rebuffered using 100K ultrafiltration columns into UB buffer (Actome) providing chemically defined conditions, finally PICOzyme buffer was added along with PICOzyme to activate the antibody. Briefly, the conserved N-glycolisation site, present on virtually all IgG antibodies regardless of isotype and host species, was trimmed to a defined structure, while PICOtransferase attaches an azide group, subsequently used for the attachment of the labels. Afterwards, the unbound labels were removed using PICOglue Antibody Binding Resin purification. The antibodies were rebuffered in PICOglue Antibody Storage Buffer using 100K ultrafiltration columns and stored at 4 °C. Finally, the concentration of the labeled antibody was determined by dPCR (by counts of labels). The labeling efficiency, *β*, was determined by measuring the molecular weight shifted fraction of heavy chain using the Protein 230 Kit (5067-1517, Agilent) run on Agilent’s Bioanalyzer under reducing conditions according to manufacturer’s instructions. On the electropherogram the light chain (∼ 26 ± 3 kDa), the unlabeled heavy chain (∼ 50 ± 4 kDa) (*δ*_*u*_), and the labeled heavy chain (∼ 75 ± 5 kDa) (*δ*_*l*_), are identified and the concentrations are determined. Considering the molecular stoichiometry of antibodies, *β* can be calculated as

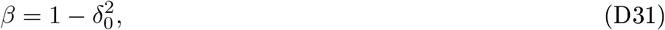

where *δ*_0_ is the unlabeled fraction of the heavy chain *δ*_*u*_*/*(*δ*_*u*_ + *δ*_*l*_). *β* is used to calculate labeling correction for the antibodies and couplex concentrations.

### D.4 Digital PCR

Generally, bold symbols are counts, light symbols are concentrations.

Droplet digital PCR (ddPCR) using Bio-Rad’s QX200 Droplet Digital PCR System was according to manufacturer’s instructions. Briefly, the ddPCR master mix were prepared using 1X ddPCR Supermix for Probes (No dUTP) (1863024, Bio-Rad), 800 nM forward and reverse primer (PICO AMC Kit, Actome), 250 nM BL and P8 probe (PICO X Probe Kit, Actome), 1 μl template and ultra pure water, up to 23 μl per sample. 20 μl master mix and 70 μl of Droplet Generation Oil for Probes (1863005, Bio-Rad)were loaded into the DG8 Cartrigdes (1864008, Bio-Rad) and emulsion generated using Droplet Generator (1864002, Bio-Rad). A C1000 Touch*™* Thermal Cycler (Bio-Rad) was used for thermal cycling of the emulsion, initial denaturation (95 °C for 10 min), followed by 40 cycles of 95 °C for 30 sec and 58 °C for 1 min, and finally 98 °C for 10 min. Droplets were analyzed using the corresponding Droplet Reader (1864003, Bio-Rad) and software. Data was exported and analyzed PICO software.

Crystal dPCR (cdPCR) using Stilla’s naica system was performed according to manufacturer’s instructions. Briefly, the master mix was prepared using 1X PerfeCTa Multiplex qPCR ToughMix (95147-01K, Quantabio), primers and probe as above, 1 μl template, 0.23 μl background dye fluorescein (100X, according to Stilla Technologies’ instructions), up to 23 μl H2O per sample. 20 μl of master mix were loaded into Sapphire Chips (Stilla Technologies). The cycling conditions were, partitioning at 1 bar overpressure, initial denaturation (95 °C, 10 min), followed by 45 cycles of 95 °C, 30 s and 58 °C, 15 s, and finally the pressure was released. The chips were analyzed in the Prism3 reader (Stilla Technologies) with the following settings, 65 ms and 150 ms exposure time for FAM and HEX channel at 82 mm focus.

Partition based dPCR (pdPCR) using QIAGEN’s QIAcuity Digital PCR System was performed according to manufacturer’s instructions. Briefly, the master mix was prepared using 1X QIAcuity Probe PCR Kit (250101, QIAGEN), 800 nM forward and reverse primer (PICO AMC Kit, Actome), 400 nM BL and P8 probe (PICO X Probe Kit, Actome), 1 μl template, up to 42 μl ultrapure water per sample. For higher degree multiplexing (three or four colors), N6 and O7 probes were also added. Also, for some experiments LNA probes were used (FAM-iowa Black FQ, HEX-iowa Black FQ and Tex615-iowa Black RQ, iDT). 40 μl master mix were per wells of a Nanoplate 24-well (250001, QIAGEN). The cycling conditions were, standard priming profile for Nanoplates 24-well, initial denaturation (95 °C, 2 min), followed by 40 cycles 95 °C, 15 s and 58 °C, 30 s. The Nanoplates were imaged using the following conditions, green channel 500 ms exposure time, gain 6, yellow channel 400 ms exposure time, gain 6, orange channel 400 ms exposure time, gain 6, and red channel (300 ms exposure time, gain 4).

The probes correspondingly detect the P8 label with FAM dye (green, *λ* = 517 nm), BL label with HEX dye (yellow, *λ* = 559 nm) emission, N6 label with Atto550 dye (orange, *λ* = 575 nm) emission, and O7 label with TexasRed dye (red, *λ* = 559 nm) emissions, however technical implementations and naming of channels are varied with instruments.

Corresponding softwares of the dPCR instruments provide ***b***_**01**_, ***b***_**02**_, ***ξ*** counts and *m* (number of compartments) per master mix volume, these counts were further adjusted to accommodate corrections for dilution errors (*λ*-normalization), dPCR clustering bias (ABC correction) and labeling normalization (*B*-normalization), D.5. After these steps the counts were converted into concentrations of the *p* = (*b*_01_, *b*_02_, *ξ*) model.

### D.5 PICO

In the isomolar titration (IMT) PICO assay, the antigen concentration remained constant while the antibody concentration was varied (C7). IMT is applied to determine the saturation concentration of the antibodies. On the contrary, the AQ curves were generated at varying antigen concentrations, with the antibody concentration held constant. PICO assays were carried out according to the protocol of the manufacturer (PICO Amplification Core Kit, Actome). Briefly, PICO measurements were performed by incubating (lysed) samples with a mixture of PICO-labeled antibodies (antibody mix - ABX) overnight, termed binding reaction, combining 2 μl of each in LBTW. The concentration of antibodies applied were predetermined (using IMT) and for absolute quantitative measurements saturation or, if it is higher, the default concentration was applied, the default is 5x10^−10^ M. Subsequently, the binding reactions were diluted in PBS to target at least 0.15 of antibodies per dPCR compartment (Poisson mean = *λ* = 0.15) for each antibody. Digital PCR (dPCR) was conducted according to the standard dPCR protocols as described earlier. For the negative control, zero couplex control, antibodies were mixed without antigen in ABC buffer, but otherwise subjected to the same processing steps as the samples (referred to as the antibody-binding control, ABC). With 8-plex PICO, oligonucleotide labels were amplified using two distinct primer pairs. This allowed the PICO reaction to be split before dPCR and the two sets of four labels to be independently amplified and read.

The raw couplex counts were first *λ*-normalized (see below), then corrected against the signal of ABC control, which serves as the zero couplex reference for the identically processed samples correcting dPCR clustering biases, and finally the labeling efficiencies were also compensated using corresponding Δs further correcting the antibodies and couplex values. Lambda normalization corrects for processing-introduced variations of *ξ* under the assumption that replicates should share the same lambda (*λ*) values. This enables the normalization of couplex counts based on the differences in *λ* among replicates. The correction normalize couplex counts toward the replicates’ mean while the mean is not changed.

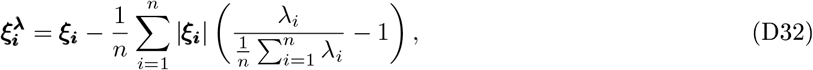

where *n* is number of replicates, 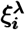 is the lambda-normalized couplex count for a given replicate and *λ*_*i*_ is the average number of antibodies per compartment for the given replicate, default both antibodies’ *λ*s are used and the calculated ***ξ***_***i***_ counts are averaged for the given replicate. To account for the ABC zero reference, all lambda normalized ***ξ*** counts are shifted by the average ABC ***ξ*** offset from zero. The ABC ***ξ*** offset indicates Poisson non-compliant clustering bias indicating not correctly clustered compartments, underestimating the number of couplexes, hence it is usually negative. Positive and large negative values of ABC ***ξ*** offsets need to treated carefully signaling experimental errors. Finally, *B*-normalization is used to correct both the counts of antibodies and the ***ξ*** count to account for unlabeled antibodies, dividing antibody counts with the respective *β*s (see B.4).

The adjusted antibody and couplex counts are transformed into concentrations (*b*_01_, *b*_02_, *ξ*) of the binding reaction, taking into account the applied dilution before the dPCR step. To calculate the AQ results, these concentrations are substituted into the *p* = (*b*_01_, *b*_02_, *ξ*) model, utilizing a surrogate *K*_*d*_ of 10 pM. To opt the appropriate solution from 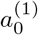 and 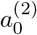, samples were measured with two known dilution factors.

### D.6 Analytical PICO

A dimer composed of BL and P8 labels (mocking couplexes) from Actome underwent a 2-fold HPLC purification process (metabion, Planegg). These dimers are detectable by both BL and P8 probes (Actome). Various quantities of dimers were combined with equimolar BL and P8 monomer labels (Actome), ensuring uniform BL and P8 counts across samples with differing dimer amounts. The synthesized dimer DNA contained synthesis errors, resulting in certain dimer molecules being non-double-amplifiable. These impurities were quantified and factored into the calculations. The samples were subjected to replication on Stilla’s naica and QIAcuity Digital PCR systems, following the previously outlined procedures. All input DNA samples were meticulously quantified, and the resulting measurement uncertainties were propagated. The limit of detection (LOD) and limit of quantification (LOQ) were determined using a York linear fit, the LOD calculation was described by Wenzl et al. [28].

### D.7 ErbB2 Western blotting

BT474 and MCF7 cells were lysed as described above. The samples were combined with 4X NuPage LDS Sample Buffer (NP0007, Invitrogen), 10 % beta-mercaptoethanol was added and heated for 5 min at 65 °C. The samples were then loaded into a NuPAGE*™* Novex 4-12 % Bis-Tris Gel (NP0321PK2, Invitrogen*™*). The gel was run for 50 min at 200 volts in NuPAGE*™* MOPS running buffer (NP0001, Invitrogen). Blotting was performed for 7-8 min using the iBlot 2 Gel Transfer Device (IB21001, Invitrogen). In the iBind*™* Flex Western Kit (SLF2002S, Invitrogen), the nitrocellulose membrane (IB23002, Invitrogen) was immersed in 1X iBind Flex Solution (SLF2020, Invitrogen) and subsequently incubated for 3-4 hours with primary antibodies against ErbB2 (trastuzumab, Roche), ErbB3 2F12 (MA5-12675, Invitrogen), S6K1 2C2 (SAB1412617, Sigma-Aldrich), 4EBP1 554 (AHO1382, Invitrogen), 4EBP1 phosphorylation T46 (MA5-27999, Invitrogen), GAPDH (600004-1-Ig, Proteintech) and secondary anti-human-HRP (SA00001-17, Proteintech) and anti-mouse-HRP (SA00001-1, Proteintech) antibodies. After rinsing the membrane with water, it was incubated with SuperSignal*™* West Pico PLUS Chemiluminescent Substrate (34580, Thermo Scientific) and chemiluminescence was detected with a imageQuant 800 Western blot imaging system (Amersham). Exposure times were as follows: 10 min for ErbB2 in BT474 cells, 10 sec for GAPDH in BT474 cells, 1 min for ErbB2 in MCF7 cells and 20 sec for GAPDH in MCF7 cells. Images were processed with Adobe Photoshop.

### D.8 Statistical analyses

Shapiro-Wilk Test was used to test the normality of the data. For multiple comparisons ANOVA was used, post-hoc comparisons were conducted using the Tukey Honestly Significant Difference (HSD) test, adjusting for multiple comparisons. For group comparisons involving non-parametric data, the Kruskal-Wallis ANOVA with Tukey’s Test was applied. For parametric data were subjected to t-Test. Non-parametric data were analyzed using Wilcoxon Signed Rank Test. To investigate the effect of drug treatment, multiple paired t-tests were utilized. Statistical analyses were executed using OriginPro 2023 (64-bit) 10.0.0.154 (Academic), complemented by appropriate data visualization tools. Significance levels were defined as follows ns (not significant) indicated a p-value > 0.05, * (significant) represented a 0.05 > p-value > 0.01. * * (highly significant) corresponded to a p-value < 0.001 or exact values were indicated.

## Acknowledgments

The authors thank Attila Karsai, Maike Smits, Bettina Moeckel, Georg Lentzen, Hardin Bolte, Martin Schwemmle, Zsolt Ruzsics and the Actome’s team for constructive discussions. This work was supported in part by contracts of Ministerium für Wirtschaft, Arbeit und Wohnungsbau Baden-Württemberg PRIMO (Az: 3-4332.62-HSG/84) and DINAMIK (Az: 7533-7-11-10-06) and Baden-Württemberg Stiftung, MONOGRAM - MET-iD55. .

## Competing interests

C.J. and P.K. have filed patents based on this work. C.J., P.K. and R.Z. are co-founders of Actome GmbH, N.G. and T.G. hold virtual shares in Actome GmbH. Qiagen is a shareholder of Actome. The remaining authors declare no competing financial interests.

## Notes

### Competing Interest Statement

C.J. and P.K. have filed patents based on this work. C.J., P.K. and R.Z. are a co-founder of Actome GmbH, N.G. and T.G. hold virtual shares in Actome GmbH. Qiagen is a shareholder of Actome. The remaining authors declare no competing financial interests.

### Summary of Updates

Revised version including line counting. No major changes.

